# NMR-measured Per-residue Electrostatic Potentials Reveal the Importance of Arginine-Lysine Balance on the Conformational Ensemble of Disordered Histone H3 Tails in Nucleosome Core Particles

**DOI:** 10.64898/2026.10.25.761679

**Authors:** Aidan B. Estelle, Vincent Schnapka, Nicolas Bolik-Coulon, Massimiliano Bonomi, Lewis E. Kay

**Author notes:** V.S. and N.B-C. contributed equally to this work.

## Abstract

Biophysical studies of chromatin have provided key insights into how the genome is regulated. Many atomic resolution structures of the fundamental building block of chromatin, the nucleosome core particle (NCP), as well as larger NCP arrays, both in isolation and in complex with effectors, or with post-translational modifications, have been elucidated. These provide in-depth pictures of key functional interactions. Considerably less information is available on the intrinsically disordered nucleosome tails that are critically important for higher order interactions in chromatin, and for transcription, DNA repair, and phase separation, and that serve as hubs for post-translational modifications. Here we focus on the tail of one of the four histones, histone-3 (H3), and measure electrostatic potentials at each of its amide proton positions in isolated NCPs via solution NMR spectroscopy. Using these potentials, in concert with molecular dynamics simulations, an ensemble of structures is calculated for WT H3 tails, as well as variants where all tail arginines are replaced by lysines (R2K) or vice versa (K2R). Our ensembles reveal that K2R tails have a higher propensity to occupy the most negatively charged region of the NCP between the DNA gyres than R2K or WT tails, while lysine enriched tails have an increased relative preference for the less negative regions near the DNA entry and exit points. The structural ensembles provide insight into differences in interactions between neighboring particles that lead to changes in diffusion and phase separation propensities between the NCP variants.

## Introduction

DNA in eukaryotic cells is packaged into chromatin whose fundamental building blocks consist of protein-DNA complexes known as nucleosome core particles (NCPs). The canonical NCP is assembled from two copies each of four histones (H2A, H2B, H3, and H4) around which 147 base pairs of supercoiled DNA are wrapped^1^ (Figure 1a). Chromatin exists in varying states of compaction, wherein DNA is accessible to nuclear proteins when loosely packed and occluded and protected when tightly packed. Such packing is dependent on intra- and inter-NCP interactions and is tightly controlled and finely tuned to regulate DNA replication^2,3^, transcription^4^, and repair^5^.

**Figure 1:**
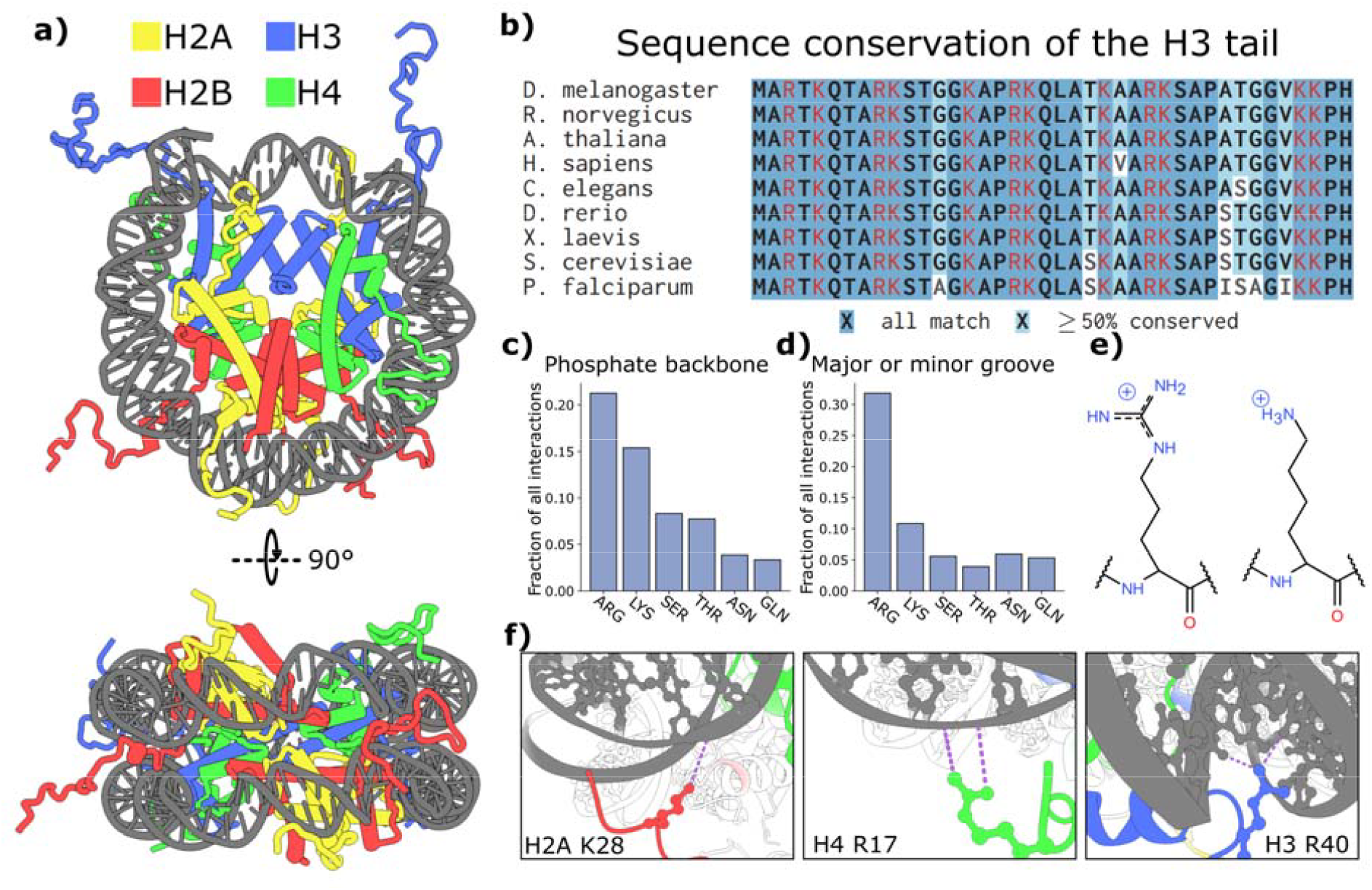
Nucleosome structure, sequence, and DNA interactions. (a) Crystal structure of the nucleosome core particle (PDB: 1KX5^*49*^). Tails have been modeled, in part based on crystal contacts; their positions are not determined confidently. (b) Sequence alignments of H3 tails across nine species with red color indicating positively charged residues. Common names D.m: fruit fly; R.n: rat; A.t: thale cress; H.s: human; C.e: roundworm/nematode; D.r: zebrafish; X.l: African clawed frog; S.c: brewer’s yeast; P.f: plasmodium (protozoan parasite). (c-d) Histogram of six most abundant residues participating in protein-DNA interactions, quantified as the fraction of total interactions involving (c) the phosphate backbone and (d) major and minor grooves of DNA. (e) Structure of arginine (left) and lysine (right) sidechains. (f) Examples of interactions between charged residues (highlighted in red, green, or blue) and nucleosomal DNA (dark grey) from available NCP structures (PDB 3UT9^*82*^ left, PDB 6L9H^*83*^ middle and right). Examples show H2A K28 interacting with DNA phosphate (left), H4 R17 interacting with DNA phosphate and sugars (middle), and H3 R40 forming hydrogen bonds with two DNA bases (right).

Much of the fine control of packing is provided by the intrinsically disordered histone tails which dynamically interact with nucleosomal DNA^6,7^. The tails extend from the N-termini of each core histone, and, additionally, from the C-terminus of H2A, and are 10-35 residues in length, each enriched in positively charged arginine and lysine residues. While the intrinsic disorder of these tails complicates their study at a structural level using most biophysical techniques, nuclear magnetic resonance (NMR) studies^8–10^ as well as molecular dynamics simulations^7,8,11^ have established that they interact with negatively charged nucleosomal DNA in a dynamic and fuzzy manner^6^. In spite of the dynamic nature of the interaction, tails protect and anchor nucleosomal DNA, regulate DNA interactions with proteins^12^, occlude DNA from nucleases^13^, and modulate DNA movement relative to the histone core of NCPs. Notably, NCPs lacking histone tails are more prone to partial unwrapping than wildtype (WT) particles^14–16^. Additionally, tail-DNA interactions can sequester the tails themselves, preventing some proteins from reading or writing post-translational modifications to them^17^. The importance of tails is further established through studies showing that tailless NCPs are unable to form liquid-liquid phase separated condensates *in vitro*^18^, and computational studies of phase separation of NCP arrays have found that tails form inter-NCP contacts that assist in condensate assembly^11^. Such nucleosome condensates serve as membraneless compartments that facilitate a variety of functions within the nucleus, including DNA repair^19,20^ and transcription^21,22^, and depend on the formation of a network of inter-NCP interactions, which are mediated in part through contacts between histone tails and DNA *in trans*^23–25^.

Many of the tail interactions, including those involving both DNA and proteins, are dominated by charge. Over a third of the amino acids in each histone tail are positively charged, and both arginine and lysine residues within NCP tails are highly conserved across species, especially for H3 (Figure 1b) and H4. Mutations and post-translational modifications that eliminate histone tail charge reduce the tails’ ability both to protect nucleosomal DNA^12,26^ and to enable NCP arrays to phase separate^23^, in a similar manner as tailless NCPs. A survey of the DNAproDB^27^ database of protein-DNA interactions highlights the importance of arginine and lysine in stabilizing nucleoprotein complexes, and by extension, the involvement of these residues in forming interactions that are required for regulating a plethora of biological activities. Arginine is notably over-represented in protein-DNA complexes, accounting for greater than 20% of the interactions formed by the DNA phosphate backbone and nearly a third of the interactions involving major or minor DNA grooves (Figures 1c-e). The high concentration of arginine can be explained, in part, by its hydrogen-bonding capability and its ability to form cation-pi interactions involving the guanidino-group^28^, as is made clear in NCP structures (Figure 1f). Additionally, the free energy penalty for desolvation is notably smaller for arginine than lysine, contributing to arginine’s more favorable association with DNA, in particular for close packing interactions within the DNA minor groove^29,30^. Recent work has shown that mutations neutralizing the charge of individual arginine residues in the H3 tail (via replacement to glutamine) have an outsized impact on the NMR-determined tail motions^31^, suggesting that arginine - DNA contacts are important for anchoring tails to the NCP. Further, mutations of arginine to lysine that preserve the tail’s net charge appear to weaken H3 interactions with DNA, as NMR-derived dynamics data indicate an increase in tail motion in these variants^18^. These substitutions also impact the properties of NCP condensates, with lysine to arginine mutations resulting in increased condensate viscosity, while the opposite set of substitutions decreases it^18^.

To understand how the high degree of lysine and arginine conservation and their specific patterning governs H3 tail interactions with nucleosomal DNA we have generated structural ensembles of the H3 tails in the context of intact NCPs, including WT, all tail arginine residues substituted by lysine (R2K), and all tail lysines replaced by arginines (K2R). This was achieved by measuring per-residue H3 tail electrostatic potentials for each variant, which we find are most negative for K2R, followed by WT, and then R2K. We use potentials as constraints to define ensembles of molecular dynamics-(MD) derived NCP structures that reproduce experimental electrostatic potential data. The resulting ensembles establish that arginine enriched tails have a larger propensity to occupy the highly negatively charged groove between DNA gyres along the side of the NCP than WT or R2K, while lysine enriched tails tend to favor less negatively charged regions of the NCP above and below the inter-gyre space and at the DNA entry and exit sites. As a result, the K2R H3 tails are more restricted in their range of conformations than WT or R2K, consistent with NMR measurements of motions for the H3 tail variants. The reported structural ensembles provide a rationale for measured differences in rates of diffusion for WT, R2K, and K2R NCPs as a function of NCP concentration, and help to explain the different phase separation propensities of these nucleosome variants.

## Methods

### Sample preparation

Histones from *Drosophila melanogaster* were expressed and purified as detailed previously^26^. Mutants of H3 (R2K and K2R) were ordered as gene blocks from Integrated DNA Technologies and cloned via Gibson assembly into a pET21 expression vector untagged. As with WT H3, cysteine 110 was mutated to a serine (C110S) in the R2K and K2R H3 sequences to prevent the formation of disulfide bonds. R2K and K2R H3 mutants were purified as described previously for WT H3^26^. H2A, H2B, and H4 were expressed in LB media without isotope labeling for all experiments in this work, while H3 WT, R2K, and K2R used in NMR measurements were expressed in D_2_O M9 minimal media supplemented with ^15^N ammonium chloride and ^2^D_7_-glucose as the sole nitrogen and carbon sources, respectively. Most of the WT assignments could be transferred to R2K and K2R from those available for the WT H3 tail^8^, as discussed below. To verify several of the assignments for the H3 K2R tail, an NCP sample was generated using H3 histones grown in D_2_O M9 minimal media with ^15^N ammonium chloride and ^13^C_6_,^2^D_7_-glucose so that high-quality triple-resonance experiments could be recorded. Unlabeled H3 WT, R2K, and K2R histones used to generate NCP samples for dynamic light scattering and condensation assays were expressed in LB media. The Widom 601 DNA sequence^32^ (147 base pairs plus 3 bases, 5’-ATC and GAT-3’, on either end from ecoRV cut sites for 153 base pairs of DNA in total) was used to wrap the histone octamers. The DNA was grown in terrific broth medium and purified as previously described^26^.

Nucleosome core particles for WT, R2K, and K2R samples were reconstituted as described elsewhere^26^. Lyophilized histones were resuspended in buffer of 20 mM Tris HCl pH 7.5, 100 mM NaCl and 6 M guanidinium chloride, mixed to final concentrations of 70, 70, 65, and 65 μM of H2A, H2B, H3 and H4, and refolded into octamers via dialysis into buffer at high salt (20 mM Tris HCl pH 7.5, 2 M NaCl, 1 mM EDTA). The slight excess of H2A and H2B over H3 and H4 prevents the formation of hexasomes, which can be difficult to separate from octamers. Octamers were further purified by size-exclusion chromatography on a 10/300 mm superdex S200 column. To prepare NCP samples, we mixed purified histone octamers with 601 DNA at a ∼1:1 ratio of 601:octamer at 6 μM concentrations of both octamer and DNA. The exact octamer:DNA ratio was fine-tuned for each sample via preparation of small-scale wrap tests over a range of calculated ratios from 0.9:1 to 1.1:1 octamer:DNA. The octamer-601 mixture was exchanged gradually into Tris buffer pH 7.5 containing 200 mM NaCl. This was accomplished by dialyzing samples in a beaker of 400 mL buffer, where both sample and dialysis buffer were at an initial concentration of 2 M NaCl. The beaker containing dialysis buffer was then drip fed, via a peristaltic pump, with Tris buffer at 200 mM NaCl at a rate of 1.5 mL/min for >18 hours, ensuring that the sample salt concentration was slowly lowered from 2 M to 200 mM over the course of many hours. Wrapped NCP samples were validated via native PAGE on 7% TBE acrylamide gels, exchanged into 20 mM MES pH 6, 1 mM NaN_3_, and stored at 4 °C until use.

### NMR spectroscopy

NMR samples were assembled from unlabeled H2A, H2B and H4, while H3 was highly deuterated and either U-^15^N labeled (electrostatics and dynamics measurements) or U-^15^N,^13^C labeled (triple resonance experiments on H3 K2R). All NMR experiments were recorded at 37 °C on samples of 60-120 μM NCP, dissolved in 20 mM MES pH 6, 1 mM NaN_3_, 3% D_2_O buffer, either with no NaCl or 100 mM NaCl. Due to the propensity of NCP samples (especially the H3 R2K NCP) to form phase separated condensates at higher salt concentrations, care was taken to exchange samples slowly into 100 mM NaCl in a centrifuge concentrator held at 37 °C, by concentrating samples to ∼500 μL and then diluting them back to 15 mL with buffer at increasing salt concentrations in 25 or 50 mM intervals up to 100 mM. All experiments were performed on a 1 GHz Bruker AVANCE NEO spectrometer equipped with an x,y,z gradient HCN cryoprobe, except for triple-resonance assignment experiments (K2R tail), which were collected on an 800 MHz Bruker AVANCE III HD spectrometer with an x,y,z gradient HCN cryoprobe. NMR spectra were processed using NMRPipe^33^ (https://www.ibbr.umd.edu/nmrpipe) and analyzed with Sparky^34^, using the peak-fitting program peakipy (https://j-brady.github.io/peakipy/) to quantify peak volumes.

#### Assignments

Assignments for the H3 R2K NCP were transferred from spectra recorded on the WT. As assignments for several peaks in the ^1^H-^15^N spectrum of the H3 K2R mutant could not be unambiguously transferred, the K2R NCP was assigned using a series of conventional triple-resonance experiments (HNCO, HN(CA)CO, HNCACB and CBCA(CO)NH)^35^.

#### Electrostatic potentials

Backbone electrostatic potentials of H3 tails were quantified via a series of ^1^H-*R*_2_ relaxation measurements recording the time decay of amide proton magnetization^36^,^37^, either in the absence (to measure baseline *R*_2_ rates), or presence of paramagnetic cosolutes, as described extensively in our prior work^9^,^38^ and in studies of others^39−43^. Negatively charged Gd-DOTA (Gd^−^) and positively charged Gd-DOTAM-BA (Gd^+^) were used in these experiments^44^ at varying concentrations depending on the sample and buffer conditions. Cosolute concentrations used for each experiment can be found in Table S1. For a pair of solvent PRE rates (one Gd^+^ and one Gd^−^), the electrostatic potential, *ϕ*_*ENS*_, can be calculated via^39^

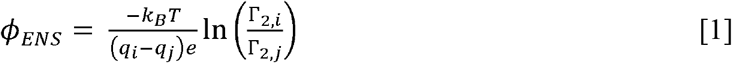

where *k*_*B*_ and *T* are Boltzmann’s constant and temperature respectively, *q*_*i*_ and *q*_*j*_ are the cosolute charges (+1 and -1 respectively in this case), *e* is electronic charge, and Γ_2_,_*i*_ and Γ_2_,_*j*_ are the solvent PREs in the presence of Gd^+^ and Gd^−^ respectively, normalized to the concentration of cosolute in each sample. To account for pipetting imprecision, cosolute concentrations were calibrated by comparing water *R*_1_ relaxation rates measured in the NMR sample against a standard curve plotting water *R*_1_ rates as a function of cosolute concentration. Standard curves were generated as described previously^9^, using the Evans method^45^,^46^ to determine cosolute concentrations via chemical shift differences of a DSS standard present in coaxial NMR tubes (5 mm and 3 mm) where the cosolute is localized to the inner tube. Potentials are reported as the average over 4-6 measurements, one for each pair of cosolutes (for example, for WT at 0 mM

NaCl, ϕ_*ENS*_ values were obtained from measurements using 0.2 mM Gd^+^ and 4 mM Gd^−^; 0.3 mM Gd^+^ and 4 mM Gd^−^; 0.2 mM Gd^+^ and 6 mM Gd^−^; 0.3 mM Gd^+^ and 6 mM Gd^−^), with uncertainties taken as the standard deviation of these values or from the propagated errors associated with the PRE rate measurements, whichever is largest..

#### J(0) measurements

*J*(0) values were obtained from a combined analysis of ^15^N R_1_ and ^1^H-^15^N two-spin {2H_*z*_N_*z*_, 2H_*z*_N_*x*_, 2H_*x*_N_*z*_, 2H_*x*_N_*x*_} relaxation rates^47^ via

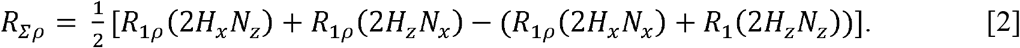

*R*_*Σρ*_ is related to spectral densities evaluated at 0, *J*(0), and *ω*_*N*_, the ^15^N Larmor frequency, *J*(*ω*_*N*_), as

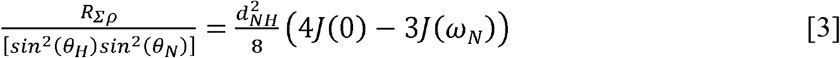

where *θ*_*H*_ and *θ*_*N*_ are given by 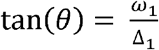, and ω_1_ and Δ_1_ are the spin-lock frequency and offset of the spin in question from the carrier frequency, respectively, for the proton (*θ*_*H*_) and nitrogen (*θ*_*N*_) spins, 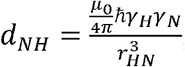, where *μ*_0_ is the vacuum permeability, *r*_*HN*_ is the distance between ^1^H and ^15^N nuclei, *ħ* is Planck’s constant divided by 2*π*, and *γ*_*H*_ and *γ*_*N*_ are the ^1^H and ^15^N gyromagnetic ratios. Note that this determination of *J*(0) assumes that *J*(*ω*_*H*_), *J*(*ω*_*H*_ ± *ω*_*N*_) ≈ 0, and therefore does not require determining *J*(*ω*_*H*_) and *J*(*ω*_*H*_ ± *ω*_*N*_) via heteronuclear NOE measurements as has been done previously^26^.

### Ensemble modeling

#### Initial models of the NCP

The Alphafold-3 (AF3) webserver^48^ was used to model the WT, R2K, and K2R NCP structures. For each sequence, 5 models were generated and the top pLDDT model was selected. The WT model closely resembled the X-ray structure of a NCP homologue sequence (PDB code 1KX5^49^), with a TM score of 0.87 and RMSD of 2.61 Å, as calculated by US-Align^50^.

#### General details of molecular dynamics simulations

All MD simulations were carried out with OpenMM v. 7.6.0^51^ using the amber99SB-ILDN force field for proteins^52^, the AMBER-parm99 parameters for DNA^53^, and the GBn2 implicit water model^54^. Implicit solvation provides a compromise between computational efficiency and accuracy, where the absence of explicit water and ions accelerates sampling of the NCP system, and the fully atomistic representation allows a more detailed model for comparison with experimental data. A cutoff of 1.6 nm was used for all non-bonded interactions, along with an implicit solvent Debye screening parameter of 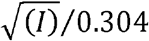, where *I* is the ionic strength (0.12 M, based on the ‘high’ salt samples studied), a solute dielectric constant of 1.0, and a solvent dielectric constant of 78.5. Simulations were performed in the NVT ensemble using a Langevin thermostat^55^ with a friction coefficient of 1.0 ps^-1^ and a time step equal to 1 fs. Bonds with hydrogen atoms were constrained with SHAKE^56^.

To preserve the structural integrity of the NCP folded domains, harmonic distance restraints were applied to specific pairs of heavy atoms belonging to all residues except those from the histone tails (H3 residues: 1-44; H4 residues: 1-26; H2A residues: 1-14 and 108-122: H2B residues: 1-35). For proteins, restraints were applied to Cβ-Cβ pairs closer than 8.0 Å in the initial model. For DNA, restraints were applied to the C2-C4, C2-C2, and C4-C4 pairs closer than 8.0 Å in the initial model, as well as to the P atom pairs closer than 14.0 Å. Additional harmonic restraints were applied to P-Cβ pairs closer than 5.0 Å in the initial model. In the case of glycine residues, the restraints listed above involved Cα carbons. All the spring constants were set to 1000.0 kJ * mol^-1^ * nm^-2^ and the equilibrium distances were extracted from the initial AF3 models.

#### Initial equilibration

A total of 15 independent NCP conformations were generated with short MD simulations initiated from the AF3 model. Each system was first equilibrated at 600 K for a total of 1 ns, after which the temperature was linearly decreased from 600 K to 300 K over 40 ns, followed by a 1 ns-long equilibration at 300 K. The final conformations were used as starting points for the production runs.

#### Enhanced sampling with simulated tempering

*Simulated tempering*^57^ was used to enhance sampling of the flexible histone tails. In this approach, a single copy of the system is simulated at a given temperature. At regular intervals, a temperature change is attempted by selecting a new temperature *T*_*j*_ among a geometric ladder of *M* =150 accessible thermodynamic states between 300 K and 800 K. Here, we used the independence sampling algorithm^58^ to select a new temperature. In this approach, the potential energy *E*_*p*_(*x*) is computed from a single conformation, *x*, and the probability that the next temperature *T*_*j*_ is selected is equal to *p*(*j*) = exp(−*E*_*p*_(*x*)/*k_B T*_*j*_ + *w*_*j*_)/*Z*, where 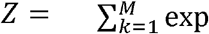(−*E*_*p*_(*x*)/*k*_*B*_ *T*_*k*_ + *w*_*k*_) and *k*_*B*_ is the Boltzmann constant. The temperature *T*_*k*_ that is selected next is determined by choosing a random number, *r*, between 0-1, and summing *p*(*j*) values from *j* = 1 (the lowest temperature, 300 K) until the sum exceeds *r*; the temperature of the rung where the sum first exceeds *r* is chosen for the next simulation. As the sampling progresses, the system will thus explore different temperatures (different rungs on the ladder), ultimately reaching high temperatures so that substantial free-energy barriers can be crossed, enabling the exploration of a broad region of conformational space. Unlike related approaches such as parallel tempering^59^, this method requires only a single replica. However, the state-dependent weights, *w*_*j*_, need to be carefully optimized to ensure rapid diffusion across the temperature ladder with approximately equal sampling for each of the *M*=150 thermodynamic states. This is achieved on-the-fly during a simulated tempering run using the Wang-Landau algorithm^60^, as implemented in OpenMM:

1. Initially, all weights are set to zero and a histogram *H* of size *M*, which measures the number of times each thermodynamic state is sampled, is initialized to 0. A weight update factor, ln(*f*), is also initialized at 1. The simulation is initiated at the lowest temperature *i* = 1.
2. The simulation is performed for 200 fs, at the end of which: *i*) the potential energy *E*_*p*_(*x*) is evaluated for the final structure; *ii*) new temperatures *T*_*j*_ are drawn with probabilities *p*(*j*); *iii*) assuming state *k* is chosen for the next iteration then the weight *w*_*k*_ of state *k* is decremented by the update factor ln(*f*) and the histogram *H*_*k*_ is incremented by 1.
3. The histogram *H* is tested for flatness. If the minimum value across all *H* entries is equal to or greater than 20% of the average, the histogram is reset to zero and the update factor is reduced by half. If a transition to a different temperature has not occurred yet, the update factor is doubled to accelerate the first transition.
4. Steps 2 and 3 are repeated for a total simulation time of 100 ns.
5. This procedure allows a first estimate of the weights. In the subsequent production simulations a different recipe for updating the weight factors is used, inspired by well-tempered metadynamics^61^. In this refinement stage, the weight update in Step 3 is replaced by ln(*f*) = *W* exp(−*H*_*i*_/Δ*T*), where *W* and Δ*T* were fixed at 1.0 and 10000, respectively. This weight refinement ensured efficient (smooth) diffusion through temperature space during the production simulation.

#### Production

For each system, 15 independent simulated tempering runs were performed starting from the 15 models obtained after the initial equilibration. Each simulation was carried out for 500 ns, resulting in an aggregated simulation time of 7.5 μs. After the first 100 ns, the coordinates of the state at 300K were saved at every Step 2, with a minimum separation of 1 ns.

#### Calculation of ϕ_PB values

Per-residue *ϕ_PB* values were calculated from our NCP ensembles via the adaptive Poisson-Boltzmann solver (APBS) program^62^ modifying protocols that have been described previously^38,44^. APBS was run using the full nonlinear Poisson-Boltzmann model and the potential grid was set as a square box of size 250 Å per side, with 481 grid points in each dimension for a grid spacing of 0.521 Å. Example APBS input files detailing all input parameters are provided in a Zenodo data deposition file (DOI: 10.5281/zenodo.22210862). APBS requires PQR formatted structures, which include atomic radii and partial charges in addition to atomic coordinates. We converted PDB files to PQR format using the pdb2pqr conversion software^63^, with protonation states calculated at pH 6 using PROPKA^64^and atomic charge and radii calculated using the AMBER force field^53^,^62^. For each amide proton *i* in each tail *n*, we computed two sums over the APBS grid points *g*,

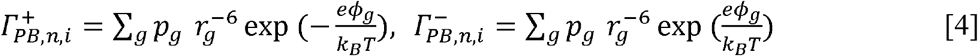

where *□*_*g*_ is the potential generated by APBS at grid point *g, r*_*g*_ is the distance from that point to the amide proton in question (amide *i*), and *p*_*g*_ is set to 1 (0) if the point is accessible (not accessible) to gadolinium cosolutes (radius 3.5 Å). Excluding constants that are assumed to cancel in the ratio^39^, 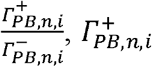, *and* 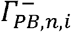 are the calculated counterparts of the solvent PREs that are measured experimentally in the presence of Gd*□* and Gd*□* cosolutes (*Γ*_2_,_+_, *Γ*_2_,_−_), respectively, for a static structure. Then the Poisson-Boltzmann potential is^39^

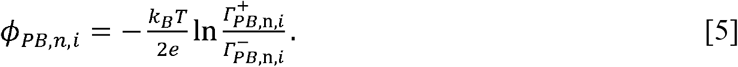

However, the measured PRE values for dynamic molecules or dynamic parts of molecules in solution, such as is the case for H3 histone tails in the context of NCPs, are ensemble averages. Therefore, for an ensemble of structures the electrostatic potential must be calculated as

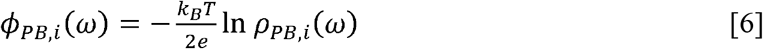

Where

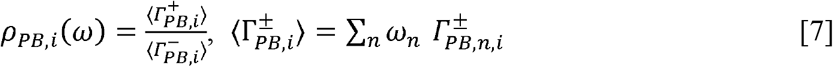

and the summation is over all *n* tails. In Eq. [7] *ρ* _*PB,I*_ (*ω*) is the calculated ratio of solvent PREs, and *ω*_*n*_ is the appropriate weighting factor for each tail (see *Maximum Entropy reweighting of ensembles*, below). Note that *ϕ*_*PB,i*_(*ω*) ≠ ∑_*n*_ *ω*_*n*_ *ϕ* _*PB,n,i*_, as the latter expression does not represent what is achieved experimentally. In total, conversion of PQR formatted structures, Poisson-Boltzmann calculations, and extraction of 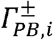values required a wall clock time of 4-6 hours per structure on 4 CPU cores, or ∼7500 hours per ensemble. To this end, calculations were run on Calcul Quebec and the Digital Research Alliance of Canada’s Narval computing cluster, with which we were able to process each ensemble in 2-4 days. Example scripts for the calculations are available in a Zenodo data deposition file (DOI: 10.5281/zenodo.22210862).

#### Maximum Entropy reweighting of ensembles

The enhanced sampling MD simulations described above generate a prior distribution,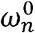, the probability of tail *n* in the ensemble of *N* tails from the *N/2* harvested NCP structures at 300 K (recall that there are two H3 tails in each NCP). Typically, it is assumed that 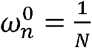that is equal probabilities for each (tail) member of the ensemble. The objective of maximum entropy reweighting is to determine a new distribution, *ω* _*n*_, that remains as close as possible to 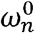while satisfying a set of *k* experimental constraints, the measured ratio of PRE rates for residue *i*,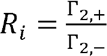 with 1≤*i* ≤ *k*. The statistical distance between the two distributions is measured by the Kullback–Leibler (KL) divergence^65^, written as a sum over tails

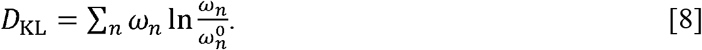

To avoid overfitting the data we follow the approach of Cesari *et al*.^66^ where the constraints are relaxed by introducing an auxiliary variable *ε* _*i*_ for each data point, representing the deviation of the calculated observable from its experimental target. As shown below, at convergence *ε* _*i*_ is the fractional deviation between the measured and calculated PRE ratios, (*R*_*i*_ − *ρ*_*PB,i*_(*ω*)) / *R*_*i*_. The *ε*_*i*_ values are treated as random variables drawn from a Gaussian prior with standard deviation *σ*_*i*_. The parameter *σ*_*i*_ encodes the level of confidence assigned to the *i*-th experimental data point. In the limit that *σ*_*i*_ → ∞ for each *i*, no confidence is placed in the data and the reweighted ensemble reduces to the prior 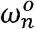; conversely, *σ*_*i*_ → 0 enforces exact matching of the *i*-th observable. Selecting appropriate values of *σ*_*i*_ is therefore critical to balance confidence in the force field, experimental measurement uncertainties, and forward model inaccuracies (see below).

As mentioned above, the experimental restraints are the set of *R*_*i*_ values and the goal is to find values of *ω*_*n*_ such that *R*_*i*_ and *ρ*_*PB,i*_(*ω*) are within experimental error. Because *ρ*_*PB,i*_(*ω*) (Eq. 7) is a nonlinear function of the weights, it cannot be used directly as a constraint in the standard maximum entropy formalism, which requires observables that are weighted averages of per-structure quantities. We therefore recast each constraint in an equivalent linear form. The condition *R*_*i*_ = *ρ*_*PB,i*_(*ω*), or equivalently, *ϕ*_*PB,i*_(*ω*) = *ϕ*_*ENS,i*_, is satisfied if and only if 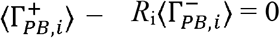, which is linear in the weights, *ω*_*n*_.

To find values of *ω_n* we begin with the full Lagrangian,

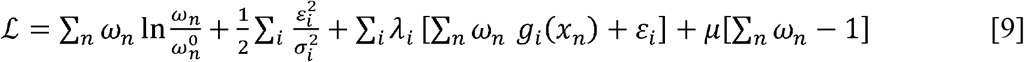

where the summations over *n* and *i* extend from 1 to *N* (all copies of NCP tails) and from 1 to *k* (the number of tail residues for which experimental PRE ratios are available), respectively. The multipliers *λ*_*i*_ and *μ* enforce the constraints and normalization, respectively, and

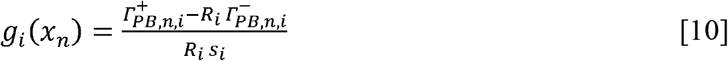

where *s*_*i*_ > 0 is a reference value of 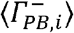 that is held fixed during each iteration cycle of the optimization process. The reweighted ensemble must then satisfy *⟨g*_*i*_*⟩ + ε*_*i*_ = 0 for all *i*, where *⟨g*_*i*_*⟩ = ∑*_*n*_ *ω*_*n*_ *g*_*i*_(*x*_*n*_), and where *ε*_*i*_ is introduced above. For *ε*_*i*_ = 0 this reduces to ⟨*g*_*i*_⟩ = 0, which is equivalent to *R*_*i*_ = *ρ*_*PB,i*_(*ω*) for any choice of *s*_*i*_. In Eq. [10] the value of *s*_*i*_ for iteration *m* is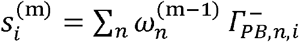, where 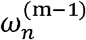 are the weights obtained from the preceding iteration and 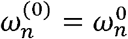. The process is repeated until *s*_*i*_ changes by less than 2% for every residue, and in our applications convergence is achieved rapidly, typically within 2 or 3 iterations. At this point 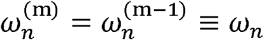, so that convergence is achieved, with 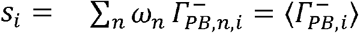.

The normalized weights *ω*_*n*_ are calculated from the requirement that

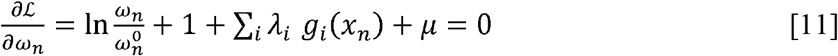

as must be the case when searching for extrema, and by imposing the normalization condition ∑_*n*_ *ω*_*n*_ = 1. Noting that because *g*_*i*_ (*x*_*n*_) is not a function of the weights, Eq. [11] can be solved in closed form

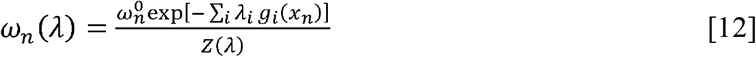

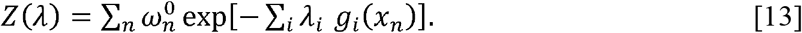

Similarly, from

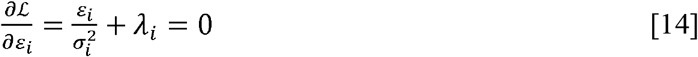

it follows that 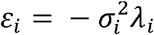. Inserting this relation and *ωn* (λ)(Eq. [12]) into *ℒ* (Eq. [9]) gives

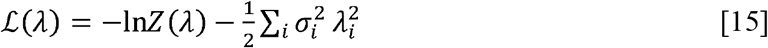

and substituting *J* = − *ℒ* gives the convex objective^67^,

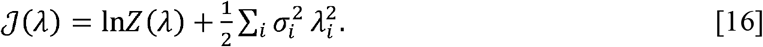

The gradient of *J*(λ) with respect to λ_i_ can be calculated

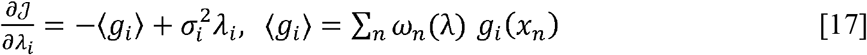

and when set to 0 gives a series of *k* equations that can be solved to get values of *λ*_*i*_, from which *ω* _*n*_ values are obtained from Eq. [12]. Note that *J*(*λ*) is convex so that a single global minimum can be found.

In absence of independent experimental data for validation, the optimal value of *σ*_*i*_ can be determined through the Kish effective sample size ratio *K*^68^, defined as

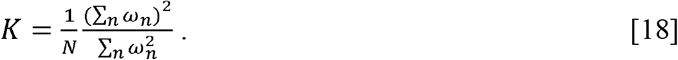

The value of *σ*_*i*_ is given by 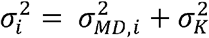, where *σ*_*MD,i*_ is an estimate of error in ⟨ *g*_*i*_ ⟩from the finite MD simulation, calculated ‘on the fly’ by the program, and *σ*_*K*_ is determined from the user set *K* value and assumed to be residue-independent. *K* ranges between 1/*N*, where only a single tail contributes to the reweighted ensemble, and 1, when all tails contribute equally. It therefore provides a direct measure of how strongly the prior distribution has been modified by the reweighting procedure. By targeting a specific value of *K*, for example in the range 0.1-0.5 ^68^, one effectively controls the trade-off between fitting the experimental data and preserving the integrity of the original MD ensemble, making *K* a practical and interpretable single free parameter for the entire reweighting protocol. For all our analyses we adjusted the Kish ratio by stepping down from 0.5 in increments of 0.025 for each set of tails (for example, WT, R2K, and K2R at a given salt concentration) until experimental data were fit to an RMSE < 3 mV. Table S2 lists the final *K* values that were selected via this protocol for each tail variant and each condition. In all cases the effective sample size of the reweighted ensemble was reasonably large (no less than 300 tails). All analyses were carried out using software developed by Robustelli and coworkers^68^ based on initial code by Cesari *et al*.^66^

Finally, the form of *g*_*i*_ (*x*_*n*_)(Eq. [10]) entering (*ℒ* Eq. [9]) is worth noting, in particular its Denominator *R*_*i*_ *s*_*i*_. This normalization ensures a similar quality of fit for each residue under a single tolerance *σ*_*K*_ independent of the value of *R*_*i*_ (i.e., the absolute value of the potential), while *s*_*i*_ factors out the raw scale of the APBS calculation (grid size, box setup) that would otherwise set the magnitude of *g*_*i*_ (*x*_*n*_). It is straightforward to show that once convergence is achieved such that 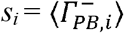, as described above, 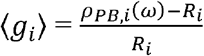, so that ⟨ *g* _*i*_⟩ reports the fractional deviation between experimental and calculated PRE ratios. Further, in this limit 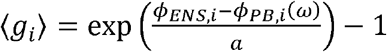, where 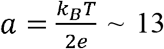 mV. In the limit where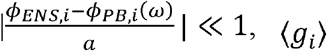, reduces to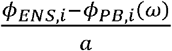.

### Visualization of structural ensembles

Density maps of tail backbones (that include N, Cα, and CO heavy atoms) were calculated by first making a 3-dimensional grid large enough to encompass backbone atoms for every H3 tail in the three NCP ensembles. The grid was comprised of voxels 2 Å in size in each dimension. Importantly for comparing density maps, the same grid size and spacing was used for all maps.

For each grid voxel *v*_*i*_, the tail probability density *p* (*v*_*i*_) for a given variant was calculated as

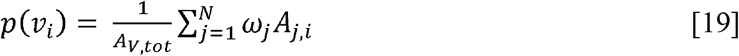

Where *ω*_*j*_ is the weight for tail *j* within the ensemble, *A*_*j,i*_ is the number of backbone heavy atoms in tail *j* within voxel *i*, and *A*_*V,tot*_ is the total number of backbone heavy atoms within tail *j* (108 for the H3 tail residues 1-36). Normalizing to *A*_*V,tot*_ ensures that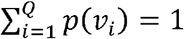, where *Q* is the number of voxels. Grids were additionally smoothed with a 3-dimensional Gaussian filter with a *σ* of 2 Å. Maps of the differences between tail densities were simply generated by subtracting one grid from another (grid 1 – grid 2), creating a new density grid where positive *p*(*v*_*i*_) values correspond to a voxel favored by grid 1, while negative values are favored by grid 2. Example scripts for calculating density grids from a set of weighted ensembles are available in the Zenodo deposition (DOI: 10.5281/zenodo.22210862).

#### Dynamic light scattering

Dynamic light scattering measurements on NCPs were collected on a Wyatt Dynapro plate reader III, in a buffer of 20 mM MES, pH 6, 100 mM NaCl, and 1 mM NaN_3_, 25°C. Measurements were carried out over a range of NCP concentrations between 5 and 40 *μ*M, and hydrodynamic radii (*R*_*h*_) for samples calculated using Wyatt’s DYNAMICS software. Reported values are the average of measurements taken in triplicate, with the standard deviation of the calculated *R*_*h*_ values set to the measurement uncertainty. To calculate 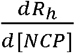 the DLS data measured for the WT, R2K, and K2R NCPs were fit to a linear model where each NCP profile shares a common *y* intercept (*R*_*h*_ at 0 *μ*M) but has a unique slope.

#### Phase Separation

To generate liquid-liquid phase separated samples of WT, R2K, and K2R NCPs, samples were initially concentrated to 350 μM (NCP) in 20 mM MES pH 6, 1 mM NaN_3_, and 1 mM EDTA and rapidly mixed with buffer containing 350 mM NaCl to a final NCP concentration of 200 μM and salt concentration of 150 mM in a total volume of 100 μL. At this point the WT, R2K, and K2R NCP samples rapidly formed liquid-liquid phase separated condensates at room temperature. Samples were allowed to equilibrate at 25 °C for 3 hours, during which time a clear separation of dilute and condensed phases formed within the sample. Aliquots of the dilute phase were extracted from the samples, diluted 100-fold into 6 M GdnCl, and quantified by measuring UV absorbance of nucleosomal DNA at 260 nm using an extinction coefficient of 0.02 (μg/mL)^-1^ cm^-1^. Each sample was measured in triplicate, with 2 hours of incubation at 25 °C between aliquots to allow samples to re-equilibrate following disruption from pipetting.

## Results

We prepared nucleosome core particles assembled from *D. melanogaster* core histones and Widom 601 DNA^32^ with H3 tails that were WT, R2K, or K2R (see Methods). As the WT H3 tail contains seven lysine and four arginine residues, the R2K and K2R variants consist of four and seven mutations, respectively (Figure S1). ^1^H-^15^N HSQC spectra recorded on NCP samples where H3 is uniformly ^15^N labeled and highly deuterated, with the remaining histones non-isotopically labelled, revealed only minor chemical shift perturbations between the variants, reflecting their intrinsically disordered nature (Figure 2a).

**Figure 2:**
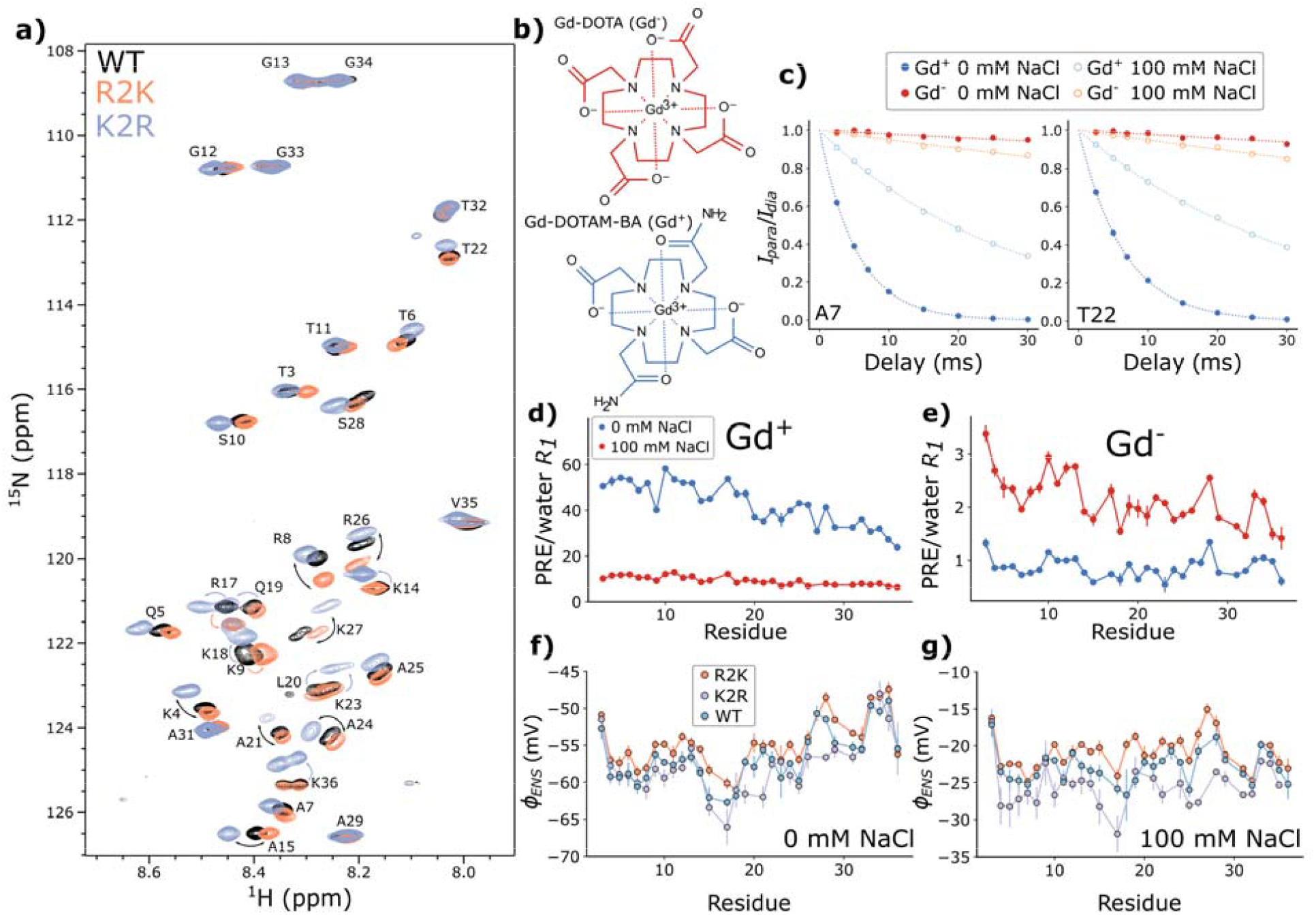
Electrostatic potentials of H3 tail variants. (a) Superposition of ^1^H-^15^N HSQC spectra of NCPs with ^2^H,^15^N-labeled H3 WT (black), K2R (lavender) and R2K (orange) tails. Assignments are shown and chemical shift perturbations are labeled with arrows. (b) Structure of Gd-DOTA (Gd^-^) and Gd-DOTAM-BA (Gd^+^), the paramagnetic cosolutes used in this study. (c) Representative plots of *I*_*para*_/*I*_*dia*_ for A7 and T22 of the WT H3 tail recorded in 0 mM or 100 mM NaCl. Rates are normalized to a cosolute concentration of 1 mM for the purpose of comparison, whereas measurements are collected at different cosolute concentrations dependent on each sample. (d,e) Paramagnetic relaxation enhancements (PREs) along the H3 tail sequence for the WT NCP at both low (blue) and high (red) salt, and for both Gd^+^ (d) and Gd^-^ (e). PRE values are normalized to measured water *R*_*1*_ rates to account for variations in cosolute concentrations between samples^*9*^. Uncertainties are based on errors obtained from *R*_*2*_ measurements and then propagated through the calculation of PREs from these rates. (f,g) Per-residue electrostatic potentials () for each H3 tail at 0 mM (f) and 100 mM NaCl (g). WT is shown in blue, K2R in lavender, and R2K in orange. Plotted values (circles) are the average of all calculated potentials using different concentrations of cosolutes (Table S1), with error bars set to the standard deviation in values, with each potential calculated from a given pair of cosolute concentrations.

### Electrostatic Potentials

We have shown in previous studies of NCPs that the measurement of per-residue electrostatic potentials (*ϕ*_*ENS*_) provides valuable insights into interactions between histone tails and nucleosomal DNA^9,38^, with negative *ϕ*_*ENS*_ values obtained for each histone amide proton, despite the large numbers of positively charged residues (arginine and lysine) in each tail. As before, here we quantified electrostatic potentials for H3 tail residues via measurement of ^1^H paramagnetic relaxation enhancement (PRE) rates in the presence of paramagnetic cosolutes of different charges, but similar structures^9,39–41,69^. We have used gadolinium metal chelated to either positively or negatively charged cages (Gd^+^ and Gd^-^, Figure 2b) and the resulting measured solvent PREs for both Gd^+^ and Gd^-^ cosolutes have been recast in terms of a *ϕ*_*ENS*_ value for each backbone amide proton using a formalism described by Iwahara and coworkers^39,69^ (Eq. [1]).

Our present data are recorded on NCP samples dissolved in 20 mM MES buffer, as opposed to 20 mM phosphate that we have used previously^9,38^. This choice reflects our interest in evaluating how *ϕ*_*ENS*_ values change for different salt concentrations (no added sodium chloride *vs* 100 mM sodium chloride), and, hence, how interactions between the H3 tail and DNA are affected by salt, and the lower ionic strength buffer (20 mM for MES vs 35 mM for phosphate at pH 6 ^9^) ensures that larger differences between the two salt conditions will be obtained. Figure 2c shows decay profiles,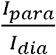, for A7 and T22, where *I*_*para*_ and *I*_*dia*_ are the corresponding peak intensities measured in spectra recorded in the presence and absence of cosolutes, respectively (separate decay profiles for *I*_*para*_ and *I*_*dia*_ are in Figure S2). This ratio eliminates the contribution from the intrinsic relaxation of the proton in question, so that the decay is due to the PRE effect exclusively. Here we have normalized intensities to those that would have been obtained with equal cosolute concentrations of 1 mM for ease of comparison, and results from experiments recorded at sodium chloride concentrations of 0 mM and 100 mM are shown. In line with our expectations that the tails would have negative *ϕ* _*ENS*_ values, the PREs for all measured proton sites in H3 are larger with the Gd^+^ cosolute than with Gd^-^. PREs for Gd^+^ are reduced upon addition of 100 mM salt, and conversely, increased for Gd^-^ across the entire H3 tail, indicating that the electrostatic potential becomes less negative with salt, as expected (Figure 2d,e). Site-specific values of *ϕ* _*ENS*_ are illustrated in Figures 2f and 2g, under conditions of 0 mM (in the range of -50 mV to -65 mV) and 100 mM (-15 mV to -30 mV) sodium chloride, respectively. Small changes in potentials are observed for each variant, following the trend | *ϕ* _*ENS*_ (R2K)| < | *ϕ* _*ENS*_ (WT)| < | *ϕ* _*ENS*_ (K2R)| and differences between each class of NCP are on the order of 2 – 5 mV. Note that the potentials for WT H3 are approximately 10 mV more negative than values we reported previously in a phosphate buffer with higher ionic strength^9^.

### Molecular Dynamics Modeling of H3 Tails

The small but systematic changes in *ϕ*_*ENS*_ values between the three tail variants suggest changes in the interactions between the H3 tails and nucleosomal DNA in each case. To obtain insight into these changes we have generated structural ensembles of the H3 tails using an approach wherein we use NMR-derived electrostatic potentials to refine conformational ensembles derived from MD simulations. A detailed description of the MD methods used to generate the NCP ensembles is provided in the Methods section and schematized in Figure 3a. Simulations were performed for a system at an ionic strength of 120 mM, equivalent to experimental conditions at 100 mM sodium chloride and 20 mM MES, as we found simulations at higher salt improved sampling efficiency. Ensembles corresponding to the low salt condition could then be generated from simulations performed at high salt by a reweighting procedure which enforces agreement of calculated electrostatic potentials to measured *ϕ* _*ENS*_ data, as discussed in detail in Methods.

**Figure 3:**
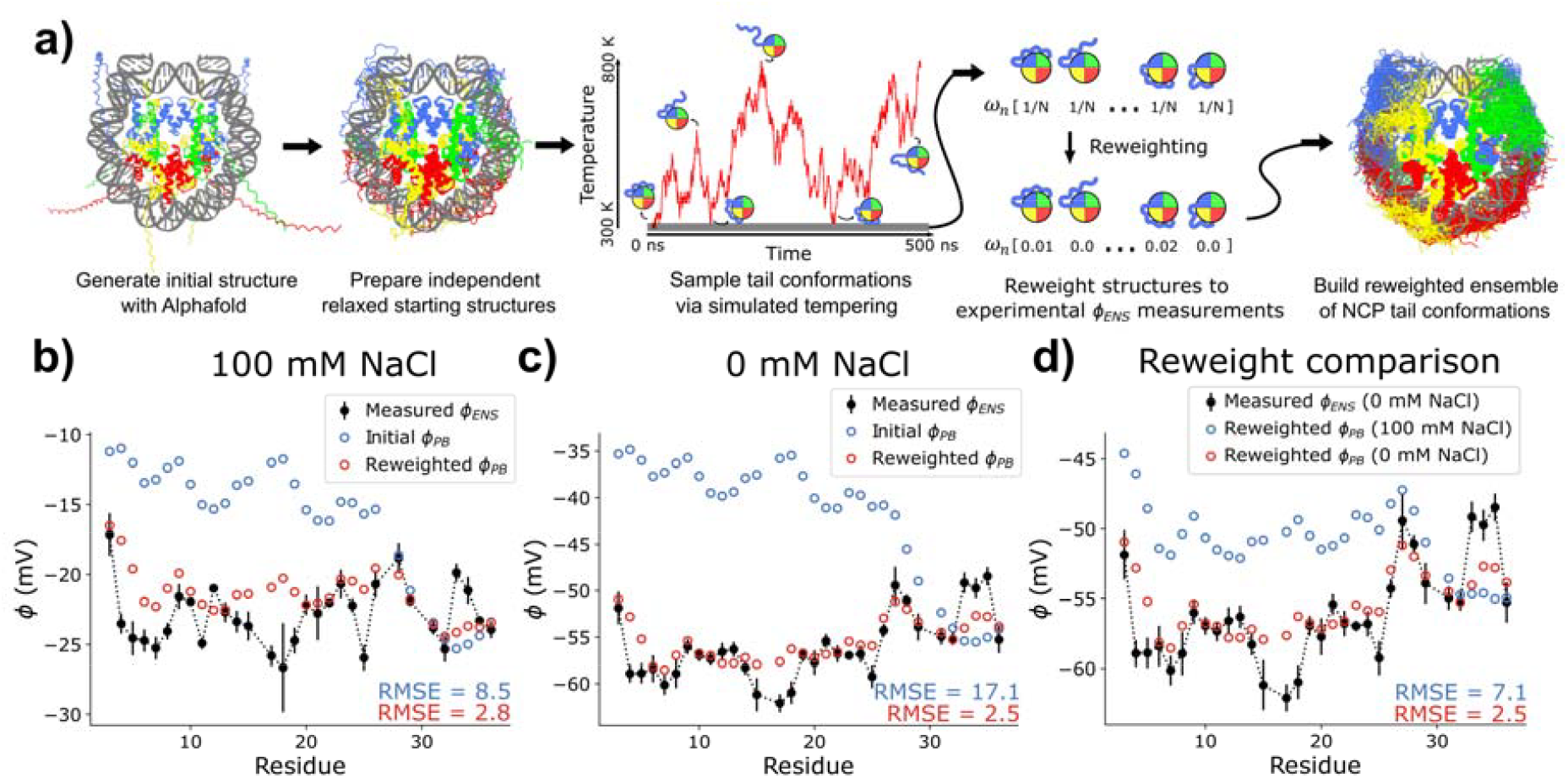
Generation and reweighting of NCP ensembles. (a) Schematic of the procedure by which ensembles of H3 tails are created. Initial structures are generated using Alphafold^48^ and then relaxed over 40 ns of molecular dynamics (from 600 K to 300 K) to the starting structures used in the simulated tempering protocol. Simulated tempering raises and lowers simulation temperatures stochastically to allow NCP tails to efficiently sample alternate conformations. From these conformations an ensemble is prepared from structures at low temperature (300 K). This initial ensemble consists of structures with equal weights, ω_*n*_, which are then reweighted to measured values, to form a final reweighted ensemble. (b,c) Effect of reweighting on ensemble potentials at 100 mM (b) and 0 mM (c) NaCl for the WT ensemble. Unweighted ensemble potentials (blue) do not match experimental data (black), while much better agreement is obtained after reweighting (red). Error bars for are based on experimental uncertainties as in Figure 2. (d) Comparison of 0 mM and 100 mM ensemble weights.Weights fitted to obtain agreement between calculated () and experimental () potentials for the 100 mM NaCl ensemble (blue) do not optimize agreement at 0 mM NaCl, as values calculated using these ‘high salt’ weights (blue) do not match experimental (black) or low salt weighted (red) potentials (0 mM NaCl). Error bars for are experimental uncertainties as in b,c. In (b-d) RMSE errors are averages over all tail residues.

In this manner we generated ensembles for WT, R2K, and K2R NCPs comprised of ∼1500 copies of nucleosome structures for each variant (and hence ∼3000 copies of H3 tails in total). We calculated ensemble mean Poisson-Boltzmann ‘rates’,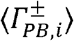, analogous to experimental PRE measurements (see Methods for details) for each residue in each tail using the Poisson-Boltzmann based program APBS^62^ via the method of Iwahara and coworkers^39^ and subsequently generated ensemble *ϕ* _*PB*_ values from the ratio of ensemble averaged Poisson-Boltzmann rates at each residue within the H3 tail. We initially sought to test for convergence of our simulations, as this is critical for obtaining meaningful structural insights in what follows, and available MD trajectories of nucleosomes include H3 tails which can diverge substantially from each other^7^. As each NCP contains two H3 tails (H3 ‘A’ and H3 ‘B’) convergence can be assessed by comparing *ϕ* _*PB*_ values calculated separately from the 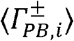rates for each. Note that while the NCP does not have perfect two-fold symmetry owing to differences in the nucleosomal DNA sequence across the dyad, the distribution of charges near each H3 tail is effectively identical, so that similar *ϕ* _*PB*_ values in H3 ‘A’ and ‘B’ would be expected if the ensemble is adequately sampled. Correlations of *ϕ* _*PB*_ values for tail residues in chains ‘A’ and ‘B’ are shown for simulations of all three variants (100 mM sodium chloride) in Figure S3a. Though a slight systematic offset can be seen in each case, differences in *ϕ* _*PB*_ between tails are small (<1 mV on average), well below the uncertainty in our experimental *ϕ* _*ENS*_ measurements. As an additional test of convergence, we split each ensemble into three randomized sub-ensembles and calculated *ϕ* _*PB*_ values at each position in the H3 tail. Differences in potentials between sub-ensembles are again within experimental uncertainty, indicating that the ensembles are adequately sampled for the purposes of our analyses (Figure S3b).

Calculated values of *ϕ* _*PB*_ (blue) are compared with experimental *ϕ* _*ENS*_ values (black) for the WT tails, for both 100 mM and 0 mM sodium chloride in Figures 3b,c, with similar comparisons for R2K and K2R shown in Figure S4. For all variants, the agreement between calculated ensemble potentials, *ϕ*_*PB*_, and measurement values is outside experimental error with systematic offsets and discrepancies as large as 5-15 mV, though the extent of variation differs across mutants and conditions (Figure 3b,c and Figure S4). This is especially true for experiments performed at 0 mM NaCl, as MD models were generated through simulations performed with a total ionic strength of 120 mM (see above), with *ϕ*_*PB*_ values subsequently calculated at 0 mM salt. We therefore chose to refine each ensemble against measured *ϕ*_*ENS*_ values. Reweighting of MD ensembles of intrinsically disordered proteins using experimental data is a powerful way to account for inaccuracies in force-fields or insufficient sampling, producing new ensembles whose weighted structures reflect the experimental data^68,70–73^. The desired outcome is a weighted ensemble of many structures that adequately describes the range of tail conformations, rather than a selection of a small number of structures that happen to agree with experimental measurements. As a result, we have used a ‘maximum entropy’ approach whereby weights for structures within the ensemble are adjusted so that the recalculated *ϕ* _*PB*_ values agree with *ϕ* _*ENS*_ measurements on a per-residue basis, while making the minimum possible number of perturbations from the initial unweighted set. Using available maximum entropy reweighting tools^68^, we reweighted each of our NCP ensembles to closely match experimental *ϕ* _*ENS*_ measurements (Figure 3b,c, compare red and black; Figure S4 for R2K and K2R). To provide an acceptable approximation of experimental error we tuned the model’s Kish ratio hyperparameter (see Methods for details) to produce a reweighted model with the largest possible Kish value while meeting the condition that reweighted *ϕ* _*PB*_ and experimental *ϕ* _*ENS*_ values agreed to within an RMSE of 3 mV (Figure 3b,c, Figure S4). As expected, the best set of weights for a given ensemble and salt concentration is not optimal for reproducing *ϕ* _*ENS*_ values measured at other conditions. For example, potentials calculated for H3 tails at 0 mM sodium chloride using weights fit to the 100 mM sodium chloride *ϕ* _*ENS*_ data are in poor agreement (systematic offset of ∼7 mV, RMSE of 7.1 mV) with*ϕ* _*ENS*_ values measured at a 0 mM sodium chloride concentration (Figure 3d, compare blue and black). Of course, the agreement is much improved with the optimally weighted ensemble (red; RMSE of 2.5 mV), reflecting changes in the structural ensemble upon addition of salt.

### Structural Models for H3 Tails

We next asked what these reweighted ensembles ‘look like’ and what unique structural features might account for the different potentials measured for the H3 tail variants examined. To get an intuitive feel for how the position of a tail in the context of the NCP might affect the underlying electrostatic potentials we initially considered each H3 tail of the 1500 ‘A’ conformations generated, calculated per-structure *ϕ* _*PB*_ values, and averaged these over the tail residues (residues 1-36) to obtain a single ‘tail specific’ value, *ϕ* _*PB,tail*_. Tails were subsequently sorted according to these *ϕ* _*PB,tail*_ values. Figure 4a shows the H3 tails corresponding to the most (red) and the least (blue) negative *ϕ* _*PB,tail*_ value for the WT ensemble superimposed on the structure of a single NCP. Interestingly, the tail with the most negative potential is inserted into the groove between the top and bottom DNA gyres (Figure 4a, red tail) which is the most negative region of a tailless NCP model (Figure 4b). In contrast, the tail with the least negative potential is positioned to one side of the NCP and partially detached from the DNA surface at its terminus (Figure 4a, blue tail). Similar relative tail positions for ensemble members with the smallest and largest *ϕ* _*PB,tail*_ values were observed for the R2K and K2R ensembles as well (Figure S5).

**Figure 4:**
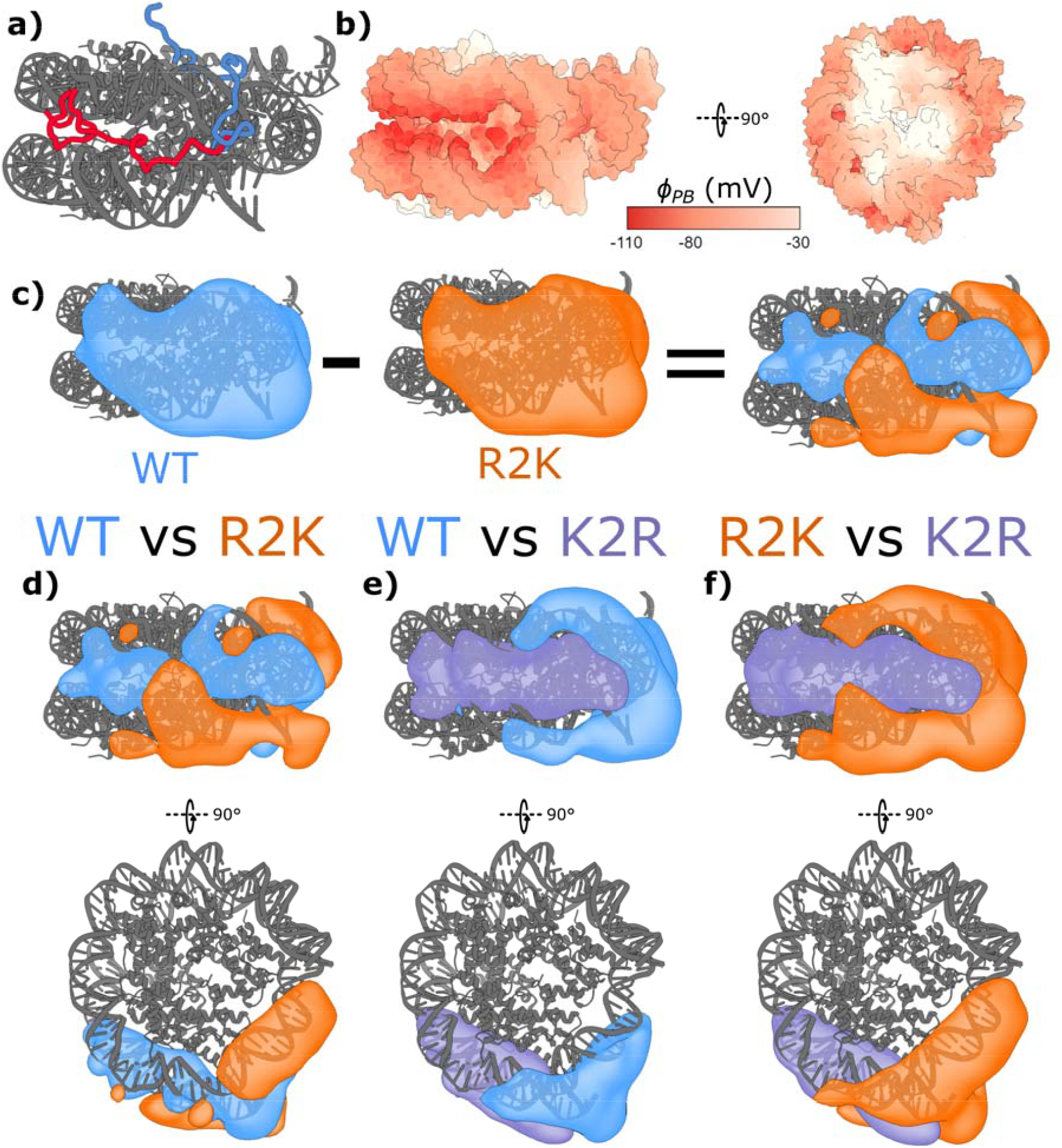
Analysis of reweighted NCP ensembles. (a) Structure of NCP from WT ensemble simulated using the 100 mM NaCl condition, showing H3 tails with the lowest (most negative, red) and the highest (least negative, blue) calculated . (b) Surface electrostatic potential of tailless NCP (PDB 1KX5^49^ with tails removed in chimeraX^84^). (c) Tail probability density maps for H3 tail ‘A’ in WT (left, blue) and R2K (middle, orange), reweighted so that and agree. Map for R2K is subtracted from the WT map to generate a map of the differences between ensembles (right) where volume favored by WT is shown in blue, and that favored by R2K in orange. (d-f) Difference density maps comparing WT *vs* R2K (d), WT *vs* K2R (e), and R2K *vs* K2R (f). Densities are colored by tail variant: blue for WT, orange for R2K, and lavender for K2R. Contour levels for map visualizations are set to 0.001% (all voxels with a density value above this threshold are shown within the contour envelope).

Interestingly, we found that the average distance from each tail amide to nucleosomal DNA was largely identical across each reweighted ensemble (Figure S6), with an average distance of ∼4.3 Å from each amide to the nearest DNA heavy atom. Thus, the tails within each reweighted ensemble are largely in contact with the DNA surface for the three H3 variants considered.

To further explore how the H3 tail conformational ensembles differ between the three tail variants, we built weighted probability density maps of H3 tail locations for each ensemble using the weights for each tail as calculated via the maximum entropy approach described in Methods. These maps are assembled as three-dimensional grids, where the weight within each voxel in the grid is determined by the occurrence of backbone heavy atoms (N, Cα, CO) from H3 tails within that space suitably normalized such that the sum of density over the entire grid totals 1. For example, a probability of 0.001 within a given grid space indicates that 0.1% of all backbone heavy atoms in the weighted ensemble are found in that grid voxel. Mapping these probability densities onto the NCP structure enables a visualization of the space that the H3 tail backbone heavy atoms occupy. As expected, the H3 tails coat the DNA surface in an area around the site where they exit the NCP core (Figure 4c). When comparing probability densities for different ensembles, we observed slight differences in the density surfaces at equivalent contouring levels (Figure 4c, compare WT (left) and R2K (middle)). To visualize this more clearly we have generated a movie (Movie S1) showing H3 tail locations for each variant as a function of probability density contouring levels, starting from high levels where there is no signal from the tails, and progressing to lower contour values. Tail positions with the highest densities emerge first in the contouring scheme. The highest probability density regions for all three tail variants localize to the inter-gyre space, as expected, since this is the most negative region of the core particle (Fig. 4b). Approximately, 15%, 21%, and 34% of the H3 tail backbones reside there for the R2K, WT, and K2R variants, respectively.

To better compare the probability densities of our H3 variants, we subtracted one grid from another, to produce a grid of differences in tail densities between two histone variants, where positive values represent space favored by one tail, and negative values by the other. In the example difference grid to the right in Figure 4c, negative density is favored by the R2K tail, and is visualized in orange, while positive density is favored by the WT tail and is visualized in blue. We built these difference maps for each of the reweighted ensembles: WT *vs* R2K (Figure 4d), WT *vs* K2R (Figure 4e), and R2K *vs* K2R (Figure 4f). Examining first the WT *vs* R2K map, we find that the WT tails have a relative preference for the highly negatively charged space along the side of the NCP, between DNA gyres, while R2K tails prefer the edges of the NCP above and below the inter-gyre space and the space around the nucleosome dyad, where the histone octamer is wrapped by a single DNA strand (Figure 4d). In the WT *vs* K2R map (Figure 4e), this pattern is reversed, with the WT now having a relative preference for the exterior space, and the K2R mutant preferentially favoring the space between the gyres. Likewise, comparing R2K and K2R shows K2R favoring the inter-gyre space, with R2K preferentially localized around the edges of the NCP surface (Figure 4f). Overall, the propensity of the tail to sit within the inter-DNA grove increases with arginine content, while the propensity of the tail to be positioned around the edge of the nucleosomal DNA and around the dyad increases with lysine content. From the difference densities illustrated in Figures 4d-f it is calculated that approximately 10-20% of the H3 tails from each histone pair localize to distinct positions on the nucleosome (for example, blue and orange densities in WT *vs* R2K, Figure 4d, correspond to approximately 10% of the tails from each ensemble, while the R2K *vs* K2R map accounts for approximately 20% of the tails). While figures show only chain ‘A’ of H3 for clarity, the same trends are found for chain ‘B’ as well (Figure S7).

Constructing probability densities and difference density maps for the ensembles reweighted to *ϕ* _*ENS*_ measurements at 0 mM sodium chloride establishes the same overall trend as observed in Figure 4 for 100 mM NaCl, indicating that the R/K-dependent variation in tail location is robust to changes in salt concentration (Figure S8). Notably, density map comparisons of unweighted ensembles show only minor differences between ensembles (Figure S9), emphasizing the importance of the reweighting process to observe nuanced, yet significant changes in tail positions.

Figures 4d-f establish that pairs of H3 tail variants have different relative preferences in their positions. By means of illustration, Movies S2 and S3 of Supporting Information show difference density plots for R2K and K2R tails, 100 mM NaCl, as a function of contouring level (side and face views of the NCP, respectively). Notably, signal from the preferred regions of the K2R tail emerges first, localized to the inter-gyre DNA region, and subsequently the regions of the R2K tails appear around the entry/exit DNA points and along the edges of the DNA. The R2K tails are noticeably more dispersed than the K2R tails. This is apparent visually from the difference maps but also reflected in the fact that the lysine enriched tails are only observed at lower contour levels, since the density probability of finding a tail heavy atom per unit voxel is smaller in this case.

We next asked whether changes in salt concentration lead to differences in weighted ensembles for the tail variants. Using the same method of mapping differences in ensemble probability densities, we compared the 100 mM sodium chloride reweighted ensembles to ensembles at 0 mM NaCl for the WT and R2K variants (Figure 5). For these ensembles we found that H3 tails from the low salt ensemble favor the groove between DNA gyres relative to the corresponding tails from the high salt ensemble that, in turn, are preferentially localized to the periphery and the dyad. This is consistent with expectations that increased salt concentrations produce a screening effect on tail-DNA interactions, enhancing the mobility of the H3 tail so that a greater area of the DNA surface can be explored relative to the case where salt is absent, with the tails relatively more constrained to regions between DNA gyres. In the case of the equivalent comparison for the K2R variant, we were unable to observe a clear and consistent difference between the 100 mM and 0 mM NaCl ensembles, as probability densities in the difference maps were low and overall trends were sensitive to changes in the Kish hyperparameter.

**Figure 5:**
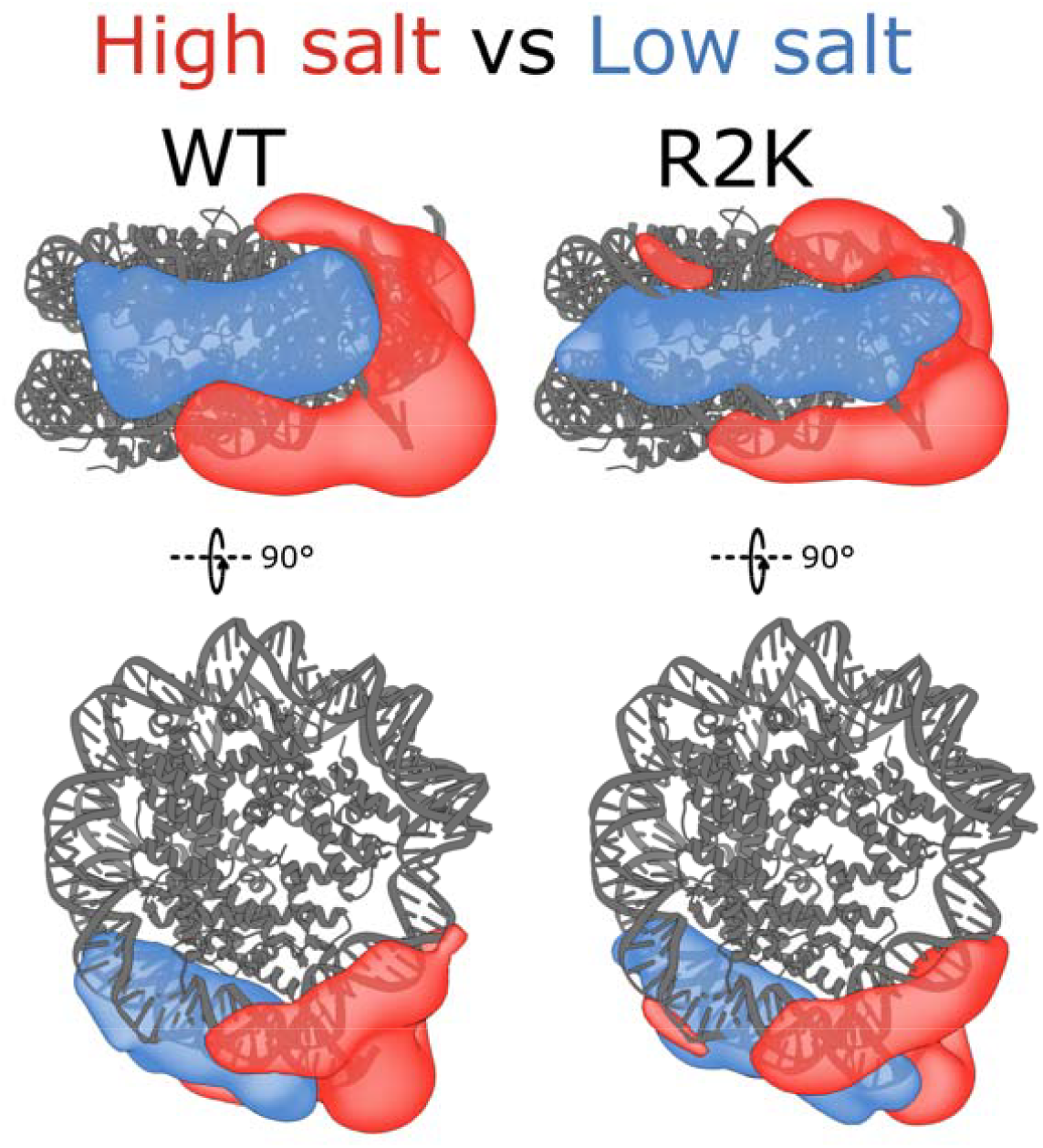
Comparison of structural ensembles reweighted to high and low salt. Difference density maps for ensembles weighted to values measured at 100 mM NaCl *vs* ensembles weighted to potentials at 0 mM NaCl. Regions favored at 100 mM are shown in red, while regions favored at 0 mM are shown in blue. Comparison is shown for WT (left) and R2K (right) ensembles. Contour levels for map visualizations are set to 0.001%. The comparison between difference density maps recorded for the K2R ensemble at high and low salt is not shown, as the results varied with the target RMSE and Kish parameter in fits, likely a reflection of little change in the position of tails in the two salt conditions.

Figures 4 and 5 highlight that the differences in *ϕ* _*ENS*_ values of the H3 variants can be explained by differences in tail location on the nucleosome surface, rather than tail unbinding from DNA. We wondered whether this result is an artifact of the tails that were used in the reweighting process to generate the ensembles. That is, do the MD trajectories favor tails that are overly interacting with DNA in the first place? To assess this, we sought to discover whether reweighting would select only those tails close to the DNA surface when structures with higher tail-DNA distances were included within the ensemble. We ‘poisoned’ the R2K ensemble (chosen as it has the least negative average *ϕ* _*ENS*_ values) with ∼1000 NCP structures (∼2000 H3 tails) taken from the maximum temperature (800 K) of our simulated tempering run that included a mix of tails near the DNA surface and extending to those unbound from the DNA (Figure 6, left). We reweighted this combined low- and high-temperature ensemble to the measured *ϕ* _*ENS*_ values. Unsurprisingly, less negative *ϕ*_*PB,tail*_ values were calculated for tails with large average amide-DNA distances, resulting in poor fits to the experimental potentials. As such, the reweighting process assigns near-zero weights to such structures, producing a reweighted ensemble built primarily of tails in close contact with nucleosomal DNA (Figure 6, right). Notably, within the unweighted poisoned ensemble, 95% of structures are within a tail mean amide nitrogen-DNA distance of 15.9 Å, while after reweighting 95% of weighted structures fall within a mean distance of 6.0 Å (Figure 6, 95% intervals are shown as dotted lines), so that unbound tails are down-weighted from the ensemble. The distribution of H3 tails along the surface of the NCP in our calculated ensembles is, therefore, not a reflection of the fact that only proximal tails to the NCP core are provided as input, but rather that close tail-NCP interactions are necessary to satisfy the measured electrostatic potentials.

**Figure 6:**
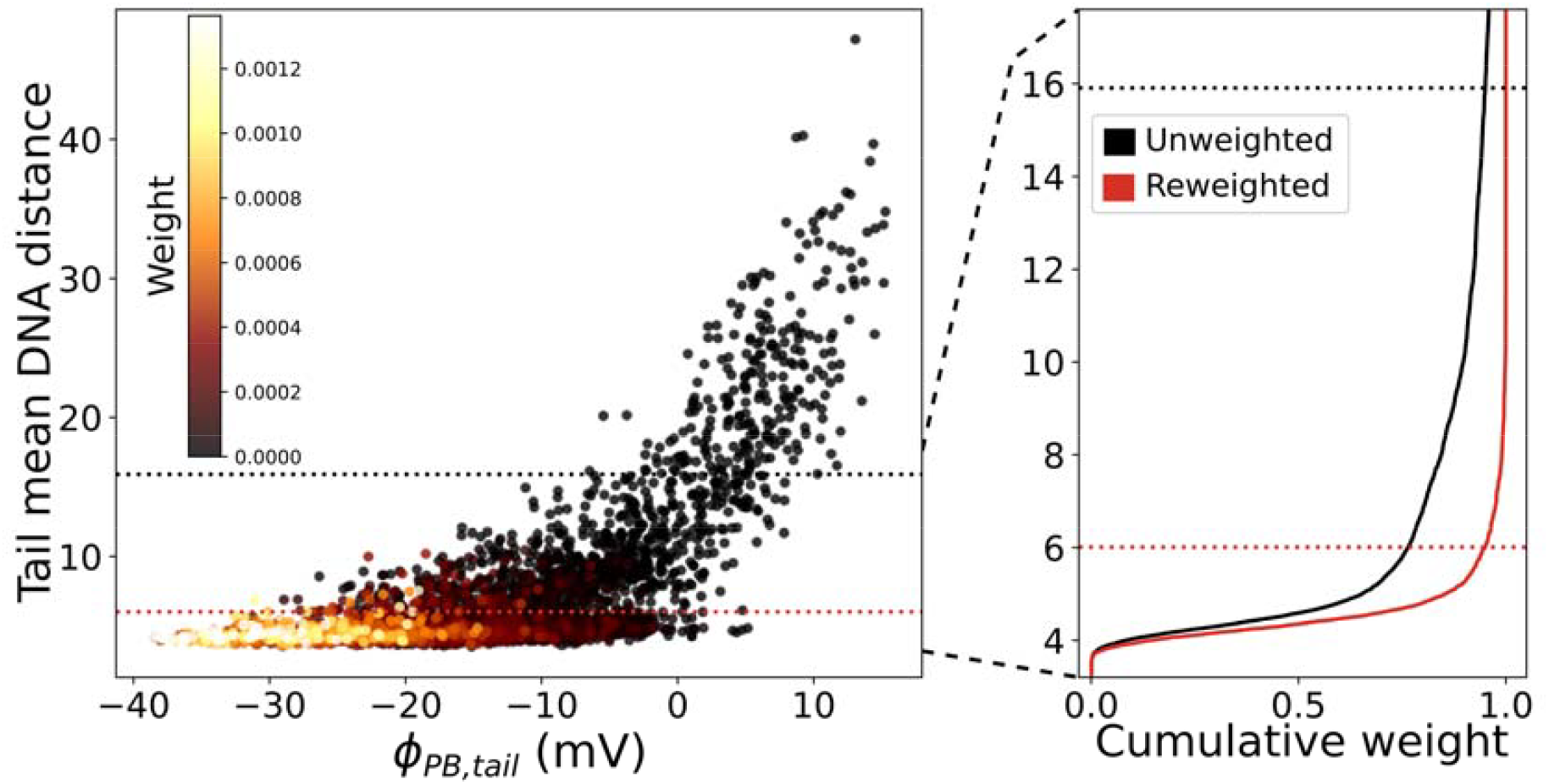
Reweighting of combined low- and high-temperature R2K ensemble establishes a clear preference for close tail-core particle distances. (left) Scatter plot of tail mean amide nitrogen-DNA distance against for each tail in the poisoned ensemble (left), consisting of 3000 H3 tails harvested at 300 K and 2000 H3 tails at 800 K. Tail mean DNA distance was calculated as the distance of a tail amide nitrogen to the closest DNA heavy atom, averaged over all residues of a given tail, and is the calculated potential averaged over all of the amides in a tail. Scatter points are colored by weight of a given tail from black to yellow (color bar inset) after reweighting to maximize agreement with values measured for the R2K NCP at 100 mM NaCl. (right) Cumulative histogram of tail mean DNA distances from 3-18 Å, adding weights from the closest to farthest distance to the DNA. Histogram for the unweighted ensemble is shown in black, and reweighted ensemble in red. Dotted lines represent the point at which 95% of the total structures are included in the cumulative weight (15.9 Å for the unweighted ensemble, 6.0 Å for the reweighted ensemble.)

### H3 Tail Fast Timescale Dynamics

Having established how exchanging lysine and arginine residues in H3 tails leads to changes in relative location preferences and interactions with nucleosomal DNA, we next tested the effect on tail motions. We recorded relaxation rates for a series of different ^1^H,^15^N two-spin coherences, as well as longitudinal order (Figure 7a), as these rates can be combined to yield chemical exchange free measures of motion on the pico-to nano-second (ps-ns) timescale^47,74^. Notably, relaxation rates increase for H3 amides in the order R2K (lowest), WT, and K2R (highest), as shown for two representative residues (A7 and T22) in Figure 7a. It is illustrative to recast the measured relaxation rates in terms of residue specific *J*(0) values, where *J*(0) is a spectral density function evaluated at 0 frequency. This is defined, in its simplest form, as 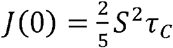, where *S*^2^is the square of an order parameter quantifying the amplitude of the motions of the H-N bond vector and *τ* _*c*_ is a residue-specific correlation time reporting the timescale of amide bond vector reorientation^75^. For intrinsically disordered regions of proteins, such as histone tails, it is difficult to separate the timescale of motions from ordering. However, qualitatively, larger *J*(0) values correspond to less dynamic and, hence more ordered, histone tails. Figures 7b,c present *J*(0) profiles as a function of residue for WT, R2K, and K2R H3 tails of NCPs measured in either 0 mM (b) or 100 mM (c) sodium chloride buffer. Values of *J*(0) vary substantially between the different tail variants at both salt concentrations examined, with *J*(0) _*R2K*_ < *J*(0) _*WT*_ <*J*(0) _*K2R*_. Thus, *J*(0) values decrease as arginine residues are replaced by lysine, consistent with an increase in ps-ns motions. Notably, *J*(0) values measured for WT and R2K samples in 100 mM NaCl buffer are lower than those at 0 mM NaCl, as salt decreases tail-DNA charge interactions leading to increases in tail flexibility (Figure 7d-f). In contrast, the salt effect on ps-ns timescale motions is small for the K2R variant, likely reflecting stronger DNA-arginine *vs* DNA-lysine interactions that persist to a greater extent in salt buffers (Figure 7f).

**Figure 7:**
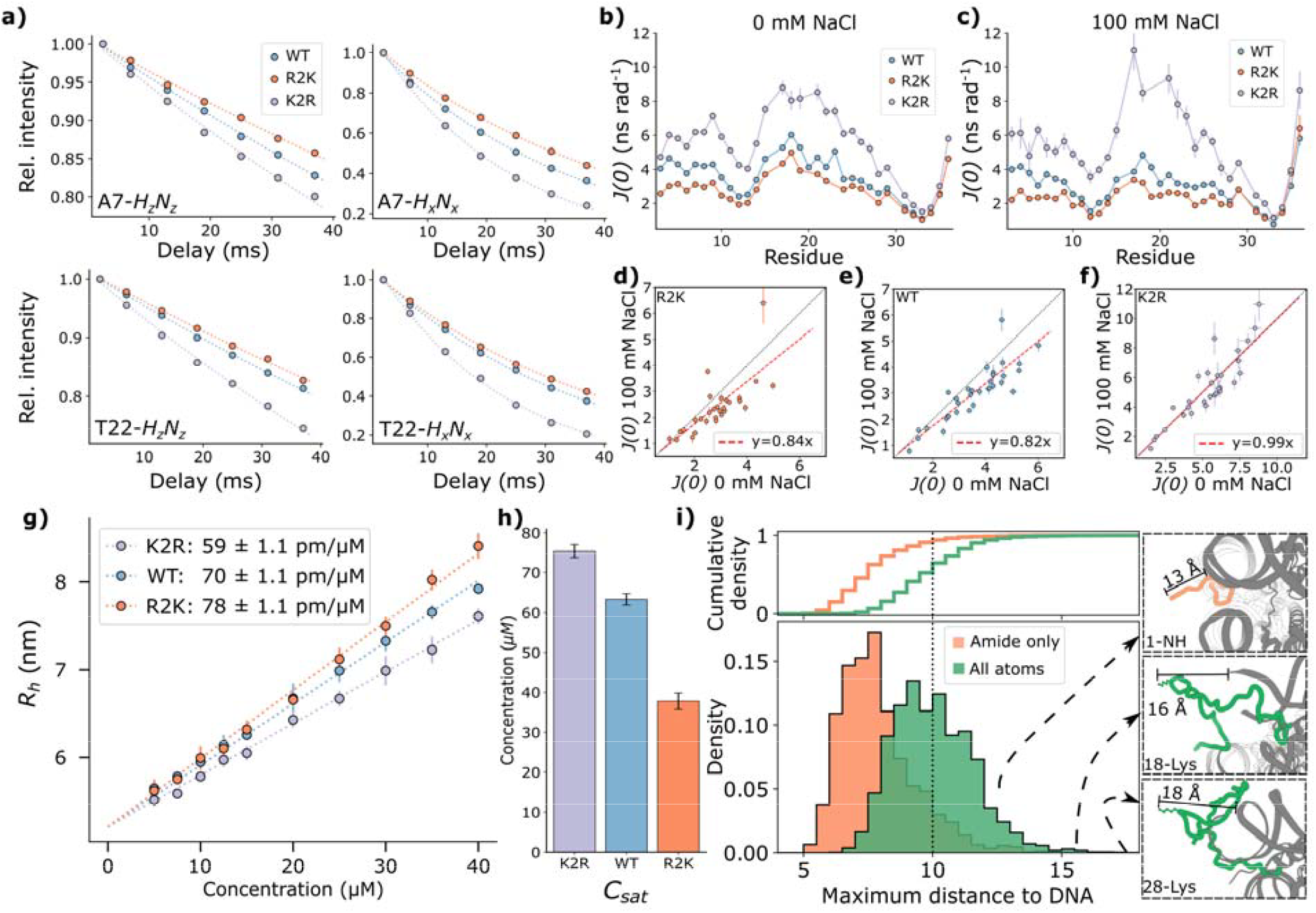
Tail dynamics, hydrodynamics, and phase separation properties of NCPs. (a) Example relaxation curves for WT (blue), R2K (orange), and K2R (lavender) H3 tails. Decays of longitudinal order (, where *A*_*i*_ is the *i*-component of *A* angular momentum) and double+zero quantum coherences () for Ala 7 (top) and Thr 22 (bottom) are shown for each H3 tail variant, along with fits of the decay curves (dotted lines). (b,c) values for WT (blue), R2K (orange) and K2R (lavender) H3 tails recorded on samples with 0 mM (b) and 100 mM (c) NaCl buffer. Uncertainty is taken from fit of relaxation decay terms and propagated to . (d-f) Linear correlation plots of values from measurements recorded at 0 mM and 100 mM NaCl for R2K (d), WT (e), and K2R (f) NCPs. Red dotted lines show fits to *y* = *mx*, with the slope, *m*, of each fit shown. (g) DLS-derived hydrodynamic radii () measurements for WT (blue), R2K (orange), and K2R (lavender) H3 tails. Dotted line shows fits to a global model with shared at 0 M NCP concentration (5.21 +/-0.02 nm). Fitted lines have a slope of which differs for each variant; values are reported in the figure inset. Errors are taken from standard deviations based on triplicate measurements. (h) Bar plot of concentration values (y-axis) measured on NCP samples with WT (blue), R2K (orange), and K2R (lavender) tails in a buffer of 20 mM MES pH 6, 150 mM NaCl at 25°C. Errors are taken from the standard deviations based on triplicate measurements. (i) Histogram (bottom) and cumulative density (top) of maximum tail-DNA distance per tail for the R2K 100 mM NaCl reweighted ensemble. Maximum distances are obtained by calculating distances for each amide nitrogen of a tail (orange) or all heavy atoms within the tail (green) to the closest DNA heavy atom, with the maximum distance chosen. Density values correspond to the fraction of tails at a given maximum distance. Example structures taken from the R2K ensemble are shown to the right, labeled with distances and the region of the tail furthest from the DNA.

#### Relating H3 Tail Structural Dynamics to Function

Histone tails form contacts with nucleosomal DNA in an intra-NCP manner but they can also interact with DNA from adjacent NCPs in nucleosome arrays to regulate chromatin packing^11,23^ and with NCPs on adjacent chromatin strands to influence liquid-liquid phase separation^18,23,24^. Inter-NCP interactions can be monitored by dynamic light scattering (DLS) measurements on samples as a function of concentration, as hydrodynamic radii (*R*_*h*_) that depend on the populations and sizes of particles can be extracted from DLS data. Figure 7g plots *R*_*h*_ *vs* NCP concentration for particles with WT, R2K, and K2R H3 tail variants, showing distinct profiles in each case. Measurements were performed at 100 mM sodium chloride, as we have shown previously that at low ionic strength interpretation of DLS data is complicated by non-idealities associated with highly charged particles^76^. Notably,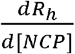values increase in the order: K2R (59 ± 1.1 pm/μM) < WT (70 ± 1.1 pm/μM) < R2K (78 ± 1.1 pm/μM). This is opposite to both the strength of intra-NCP H3 contacts probed by electrostatic measurements and the propensity for H3 tails to sit within the inter-DNA groove as observed in reweighted MD ensembles. Thus, as intra-NCP tail interactions are weakened, correlating with reduction of numbers of arginine residues in the H3 tail, inter-NCP contacts increase in magnitude.

How might this affect the propensities of the NCP variants to phase separate? To address this question, we prepared NCP condensates in buffer at 150 mM NaCl, which we found readily induces phase separation of concentrated (200 μM) NCP samples at 25°C, and measured the NCP concentration at which phase separation commences,*C*_*sat*_, for each sample (Figure 7h).

Values of *C*_*sat*_ are used as reporters of phase separation propensity, with lower concentrations consistent with higher tendencies to form condensates^77^. Measured *C*_*sat*_ values decrease in the order *C*_*sat,K2R*_ (75±1.7 μM),*C*_*sat,WT*_ (63±1.4 μM),*C*_*sat,R2K*_ (38±2.0 μM). Notably, H3 variants with more inter-NCP contacts (higher 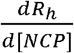values) were found to phase separate at lower NCP concentrations.

The results of the hydrodynamic and phase separation measurements indicate that inter-NCP contacts are formed, most likely very transiently. Although our structural ensembles indicate proximity between H3 tails and the NCP core, we wondered if, at least for a small fraction of the tails, some tail-core distances might be significantly larger than average to facilitate intermolecular contacts involving tails of at least one of the participating NCPs. Figure 7i shows a histogram of the maximum distance from a tail amide nitrogen to the closest DNA heavy atom for each tail from the R2K reweighted ensemble at 100 mM NaCl (orange). Distances greater than 10 Å were calculated for approximately 10% of the H3 tails, indicating that specific sites on a given tail backbone may reach out from the nucleosome core (predominantly the N-terminus of the tail), even as the remainder of the tail is in close contact with DNA. We have also calculated maximum distances for *any* heavy atom of a tail to the closest heavy atom on DNA, (Figure 7i, green histogram). A similar profile as for the backbone nitrogen atoms is obtained, shifted by approximately 2.5 Å to larger distances, where now on the order of 50% of the H3 tails have a heavy atom greater than 10 Å from the DNA. It is worth noting, however, that the positions of sidechain heavy atoms are not constrained via experimental parameters (only via the MD force field), unlike the backbone where measured electrostatic potentials are satisfied by the ensemble, so that this profile should be interpreted qualitatively.

## Conclusions

The disordered histone tails of nucleosome core particles are essential modulators of DNA accessibility and stability^12,13,15,16^, chromatin packing^11,23^, transcription^4,21,22^, DNA repair^5,20^, and phase separation^23,24^ and play important roles in the recognition of effectors to specific sites on chromatin^17,78^. This modulation occurs both indirectly through post-translational modifications^78^ and directly through interactions that reflect the biophysical properties of the tails themselves^6,7^, with charge interactions between histone tails and nucleosomal DNA critical to these processes. Though it is known that histone tails interact dynamically with DNA, experimentally derived structural ensembles of these tails and the atomic details of how the structural features of such ensembles impact chromatin behavior remain to be elucidated. Notably, histone tail sequences are highly conserved across species, including positions of arginine and lysine residues.

Here we report structural ensembles of WT H3 tails, as well as H3 tail variants where all arginine residues are replaced by lysines or vice versa (Figure 4, Movies S1 – S3). Ensembles were calculated from enhanced sampling MD trajectories where the structures generated were subsequently reweighted to achieve agreement with NMR-derived measurements of near surface electrostatic potentials of backbone amide protons. Large differences in overall *ϕ*_*ENS*_ values were observed when experiments were recorded under conditions of 0 mM NaCl (Figure 2f, 8a-top, y-axes) or 100 mM NaCl (Figure 2g, 8a-bottom, y-axes), reflecting the screening effect of salt. Notably, systematic differences in *ϕ*_*ENS*_ of a few mVs per residue were measured between the WT, K2R, and R2K H3 tail variants, leading to differences in ensembles derived for the different tail classes. While tails of all classes are proximal to the nucleosomal DNA, the R2K tails have preferences for occupying regions close to the DNA entry/exit sites and around the edges of the NCP relative to WT or K2R tails, while K2R tails localize more strongly within the highly negatively charged region between the DNA gyres (Figure 4, Movies S2 and S3). The WT H3 tails, with both arginine and lysine residues, unsurprisingly are more strongly localized between the gyres than R2K but less so than K2R tails. Additionally, a comparison of tail probability density maps suggests that tails explore a greater range of the NCP surface as the lysine content increases (Figure 4, Movies S2 and S3).

The structural analysis of the H3 tail positions is supported by ps-ns timescale dynamics measurements (Figure 7). Notably, decreased motion is observed for backbone amides for the variant where lysine residues are replaced by arginine. This effect was noted previously by Morrison and coworkers^18^ but the results here are more extensive as NCPs were reconstituted with perdeuterated H3 histones and spectra were recorded at 1 GHz, providing increased sensitivity and resolution over datasets measured on fully protonated samples at lower magnetic fields. The decreased dynamics of the arginine enriched H3 tails reflect the enhanced DNA-binding capability of arginine relative to lysine, that results from a larger number of hydrogen bonds with DNA as well as favorable cation-pi interactions involving the guanidino-moeity^28^, and the smaller energetic penalty for desolvation. Thus, the K2R tails interact more strongly with nucleosomal DNA and are more restrictive in their ability to explore the nucleosome surface, consistent with the structural ensembles calculated from the measured electrostatic potentials. The stronger contacts provide an explanation for why motions of the K2R variant are less sensitive to the addition of salt than dynamics of tails from the WT or R2K NCPs (Figures 7d-f) and help explain why clear differences between the high and low salt K2R ensembles were not observed.

Taken together, differences in the derived ensembles and the tail dynamics therein for the three H3 variants considered here provide a framework for understanding how the tails contribute to intermolecular NCP interactions (Figures 7g,7h). The interactions coupling the R2K tails with the NCP core are weakened relative to their WT or K2R counterparts, as evidenced by increased mobility and larger regions of NCP surface that are explored, as well as a preference for less negative regions of the core particle relative to either WT or K2R tails. Thus, stochastic and transient fluctuations increasing tail-NCP core distances - that would occur with higher probability for R2K than for WT or K2R variants - would lead to increased inter-particle interactions for R2K tails, as observed in our hydrodynamic and phase separation studies (Figures 7g, h) and highlighted in Figure 8b. It is well established that NCPs tend to form face-to-face stacked complexes in both mono-nucleosomes and chromatin arrays^11,79^. Further, computational studies of mono-nucleosomes have indicated that side-to-side interactions between NCPs are strongly disfavored by electrostatic repulsion^80,81^, while face-to-face and side-to-face contacts are relatively favorable. The positions of the histone tails on the DNA surface, therefore, are likely to be an important factor in bridging NCP units. When the H3 tail is preferentially sequestered against the side of the nucleosome in the inter-gyre groove, as for the K2R variant (Figure 8c) relative to the WT or R2K tails, it is less likely to contact other NCPs, as the high charge density in this region would repel nearby particles. Conversely a tail that is sitting along the edge of the nucleosome, as is favored in the R2K variant, is more likely able to promote inter-NCP interactions when a pair of NCPs make transient face-to-face or face-to-side contacts.

**Figure 8:**
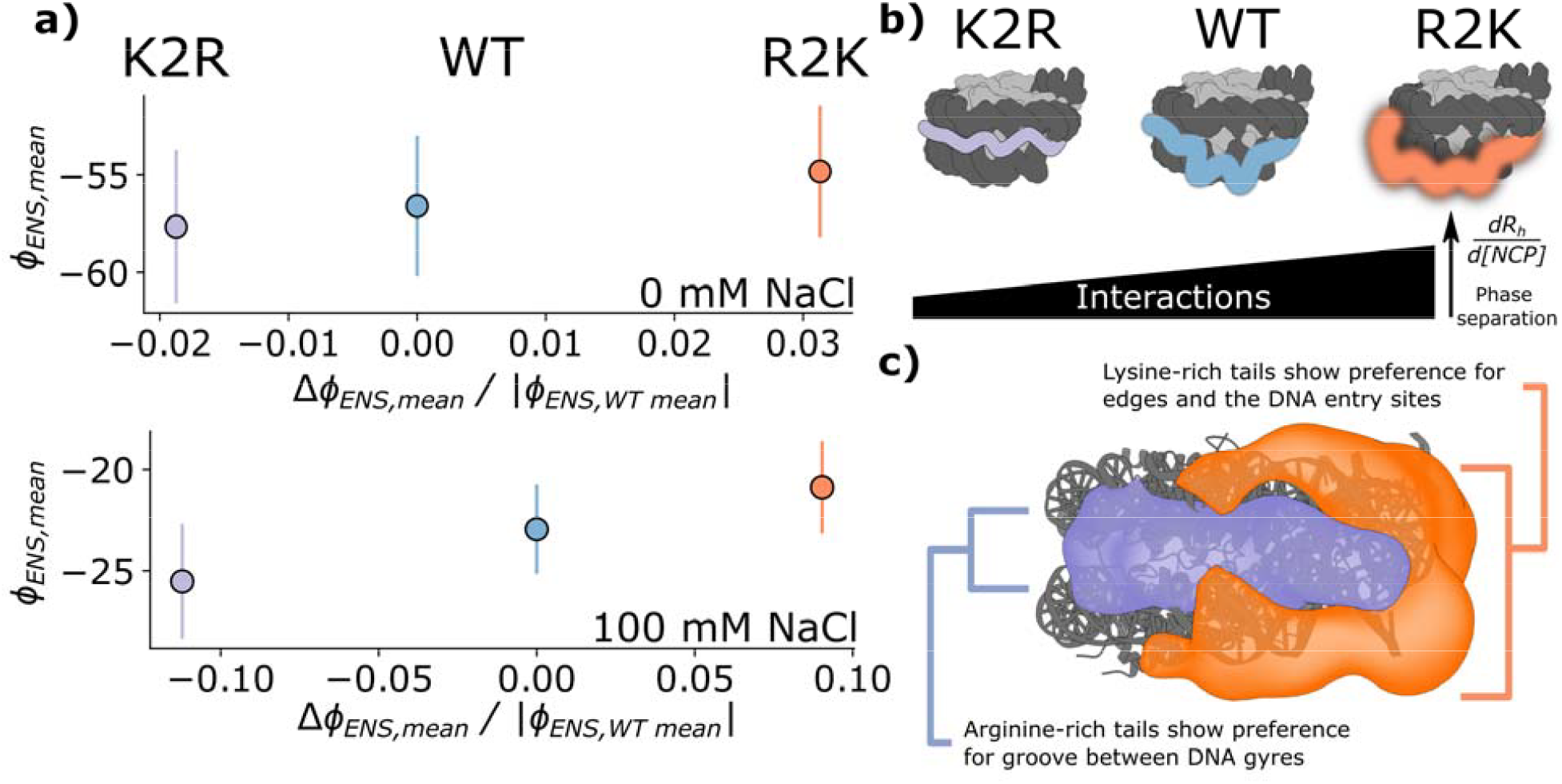
Schematic of changes in *ϕ*_*ENS*_, ensemble structures, and NCP interactions for K2R, WT, and R2K H3-tail variants. (a) Average electrostatic potential *vs* normalized potential difference 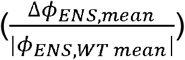for K2R (left, lavender), WT (bottom). The value Δ*ϕ*_*ENS,mean*_ is defined as *ϕ*_*ENS,j*_ *ϕ*_*ENS,WT*_, *j* ∈ (middle, blue), and R2K (right, orange) H3 tails at 0 mM NaCl (top) and 100 mM NaCl {*R2K,WT,K2R*} Vertical lines are the standard deviations of measured*ϕ*_*ENS*_ values for a given tail. (b) Schematic (top) of H3 tail structural ensembles, highlighting the relative preferences for positions of the K2R (lavender, left), WT (middle, blue), and R2K (right, orange) tails. Differences in tail ensemble structural dynamics lead to increasing inter-NCP interactions as the tail lysine content grows (bottom). (c) Diagram of R2K vs K2R difference density maps (as in Figure 4f) highlighting locations of preferential occupancy in a comparison of R2K (orange) and K2R (lavender) tails.

Here we have shown that charge-preserving mutations involving lysine and arginine residues lead to changes in H3 tail structural dynamics, with distinct relative structural preferences for each tail variant with respect to the nucleosome core particle. This, in turn, leads to changes in function through modulation of interactions involving neighboring NCPs. Previous studies focusing on charge changes through acetylation of lysine residues – one tail at a time – or PARylation of H3 suggested that these post-translational modifications are also accompanied by rearrangement of tail conformations^26,38,79^, although combined MD/electrostatic potential analyses, as performed here, were not carried out. Our study establishes that relatively subtle changes in electrostatic potential, on the order of 2-3 mV for each H3 tail residue, can be linked to changes in structural ensembles. The development of NMR methods that allow site specific measurements of electrostatic potential, even for intrinsically disordered regions of proteins such as histone tails, provides an opportunity to quantify how charge affects the structure-dynamics-function relationship in a wide array of biological macromolecules.

## Supporting Information

Supporting Information: Supporting Tables (gadolinium cosolute concentrations used in electrostatic potential measurements, list of the *K* values used for each tail variant and each condition), supporting figures (sequence of H3 variant tails, ^1^H *R*_*2*_ decays used to calculate electrostatic potentials, model convergence tests for NCP ensembles, comparison of ensemble *ϕ*_*PB*_ values before and after reweighting to experimental *ϕ*_*ENS*_ measures for each variant and buffer condition, structures of tails at minimum and maximum *ϕ*_*PB,tail*_ values for each ensemble, ensemble average distances from tail amide nitrogen to DNA, difference density maps including both H3 ‘A’ and ‘B’, difference density maps of ensembles reweighted to *ϕ*_*ENS*_ measurements at 0 mM NaCl, and comparison of difference density maps before and after reweighting), and supporting movies (density maps for R2K, WT, and K2R tails and difference density maps comparing R2K vs K2R reweighted ensembles).

## Acknowledgements

The authors are grateful to one of the reviewers for pointing out an error in our calculation of the ensemble averaged electrostatic potential in an earlier version of the manuscript. This work was supported through grants from the Canadian Institutes of Health Research (CIHR, FND-503573) and the Natural Sciences and Engineering Council of Canada (NSERC, 024-03872) to L.E.K. A.B.E. and N.B-C. acknowledge postdoctoral funding from NSERC and the Hospital for Sick Children, respectively. M.B. and V.S. were supported by the European Research Council (ERC) under the European Union’s Horizon 2020 research and innovation programme (Grant agreement No. 101086685 – bAIes). This research was enabled in part by support provided by Calcul Quebec and the Digital Research Alliance of Canada (alliancecan.ca). The linearized maximum entropy reweighting model was developed and tested with the assistance of Claude Opus 5.5, with all suggested methodology and code reviewed and tested by the authors.

## TOC Graphic

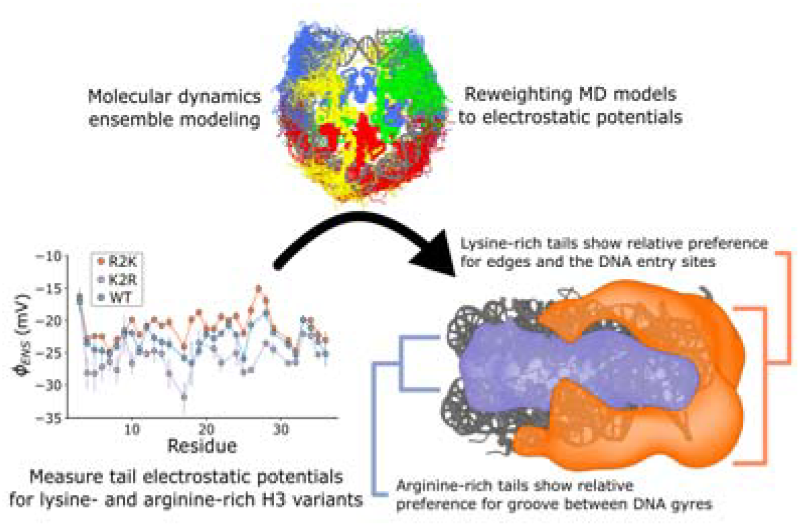

