## Supplementary tables 1-2, supplementary figures 1-9 for "NMR-measured Per-residue Electrostatic Potentials Reveal the Importance of Arginine-Lysine Balance on the Conformational Ensemble of Disordered Histone H3 Tails in Nucleosome Core Particles"

| | WT | | R2K | | K2R | |
|---|---|---|---|---|---|---|
| | 0 mM NaCl | 100 mM NaCl | 0 mM NaCl | 100 mM NaCl | 0 mM NaCl | 100 mM NaCl |
| $Gd^+$ | 0.2, 0.3 mM | 0.2, 0.3 mM | 0.2, 0.3 mM | 0.2, 0.3 mM | 0.2, 0.3 mM | 0.2, 0.3 mM |
| $Gd^-$ | 4, 6 mM | 1, 2, 3 mM | 4, 6 mM | 1, 2, 3 mM | 4, 6 mM | 4, 6 mM |

**Table S1: Gadolinium cosolute concentrations used for each electrostatic potential measurement.**

| | 100 mM NaCl | 0 mM NaCl |
|---|---|---|
| WT | 0.359 | 0.129 |
| R2K | 0.409 | 0.104 |
| K2R | 0.336 | 0.282 |

**Table S2: Kish ratios for each ensemble.** Ratios are selected via an optimization protocol initialized at target Kish = 0.5 and then stepped down in increments of 0.025 until a fit to measured $\phi_{ENS}$ with an RMSE < 3 mV is achieved. Kish ratios for 0 mM NaCl reweighted ensembles are lower as a rule due to the fact that initial ensembles are generated at 100 mM NaCl, but none optimize to a final Kish < 0.1, representing a weighted ensemble with an effective sample size of at least 300 tails.

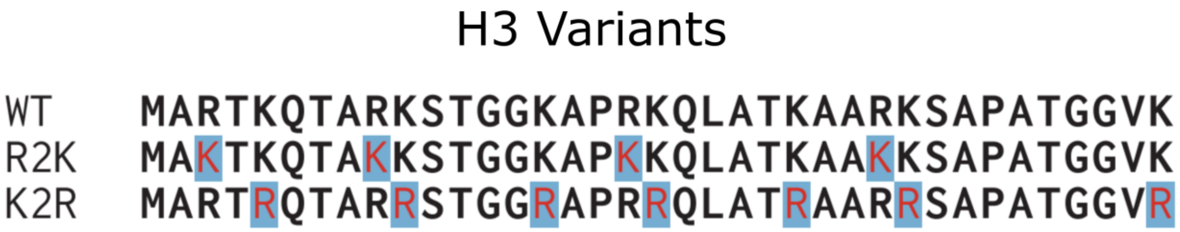


**Figure S1: Sequences of H3 variants.** WT (top), R2K (middle), and K2R (bottom) sequences of intrinsically disordered D. melanogaster H3 tails (1-36) used in this work. Mutation sites for R2K and K2R are colored in red and highlighted.

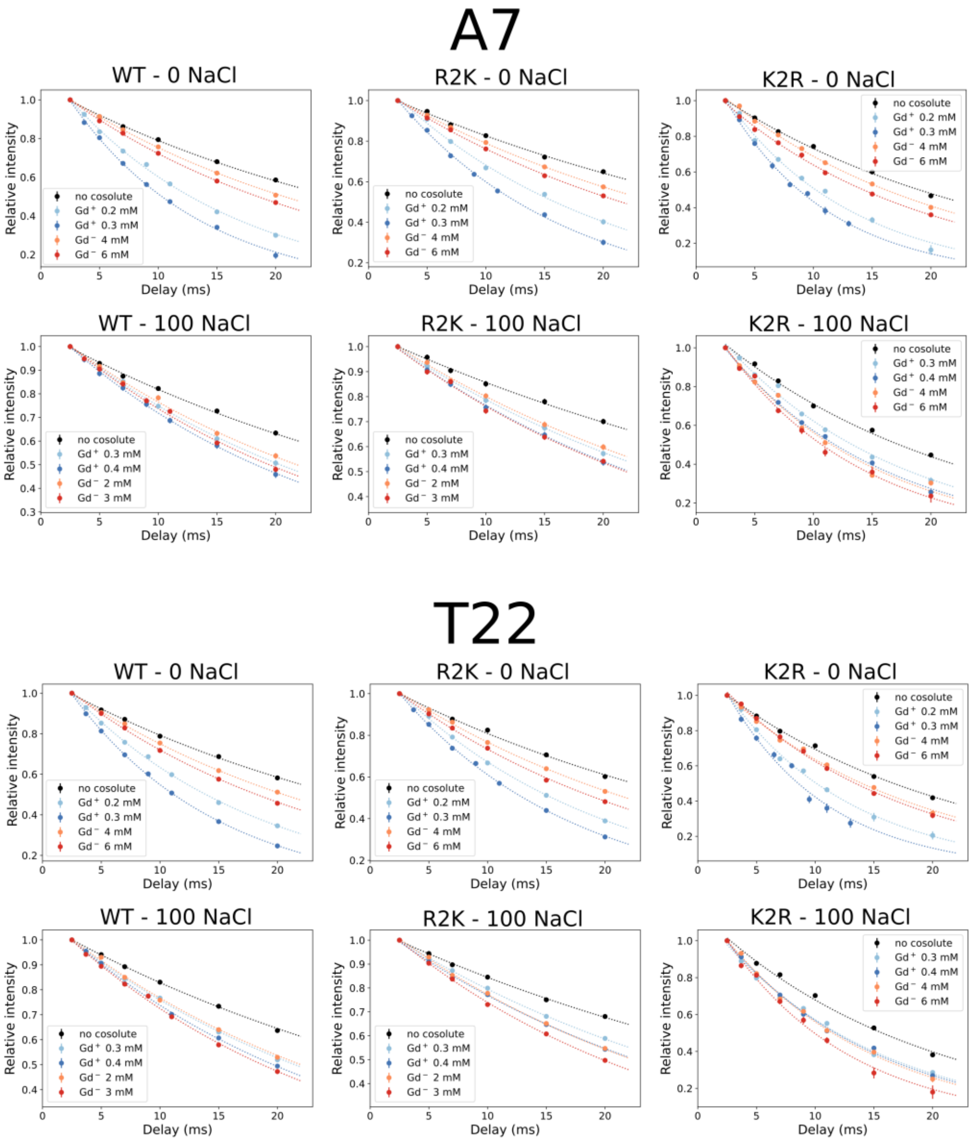


**Figure S2: Example amide-$^1$H $R_2$ decays for extraction of $\phi_{ENS}$ values.** Relative peak intensities as a function of relaxation delay (circles) and fit decay rates (dotted curves) are shown for NCP samples in the absence of cosolute (black) and in the presence of $Gd^+$ (blues) and $Gd^-$ (reds) cosolutes. Plots show measurements at both 0 mM NaCl (top) and 100 mM NaCl (bottom) for WT (left), R2K (middle), and K2R (right) NCPs for residues A7 and T22 of H3.

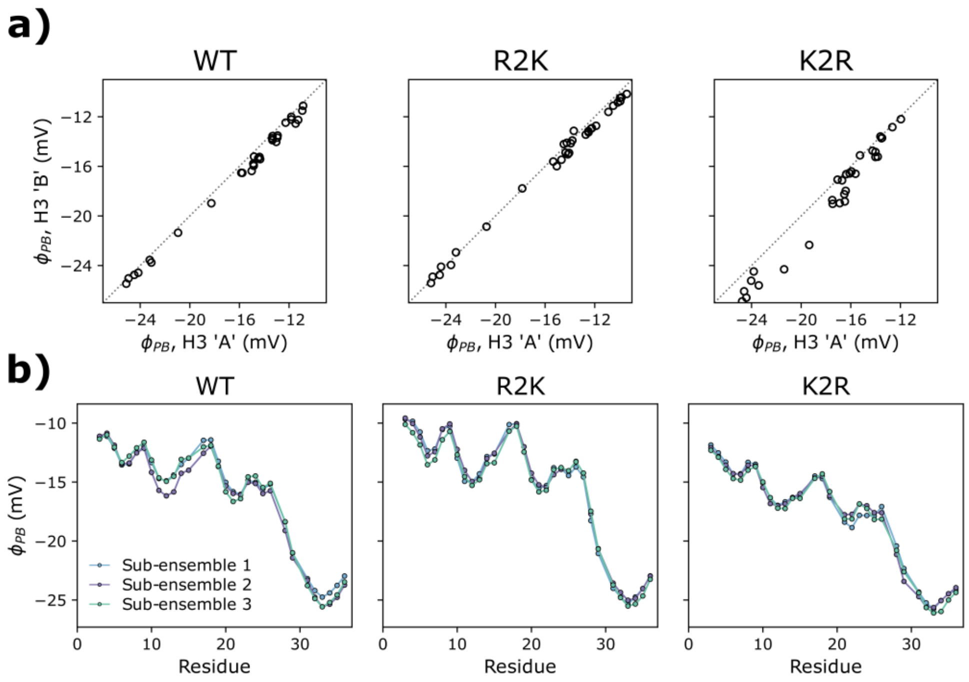


**Figure S3: Model convergence tests for NCP ensembles.** (a) Linear correlation plots of H3 tail 'A' and tail 'B' ensemble $\phi_{PB}$ values ($\phi_{PB}$) for WT (left), R2K (middle) and K2R (right) ensembles. Each point in the correlation is $\phi_{PB}$ calculated from the mean $\Gamma^{\pm}_{PB,i}$ values for a different tail residue. (b) Analysis of sub-ensembles for WT (left), R2K (middle), and K2R (right) NCPs. Ensembles are divided into 3 randomized sub-ensembles, and ensemble $\phi_{PB}$ values are calculated for each sub-ensemble.

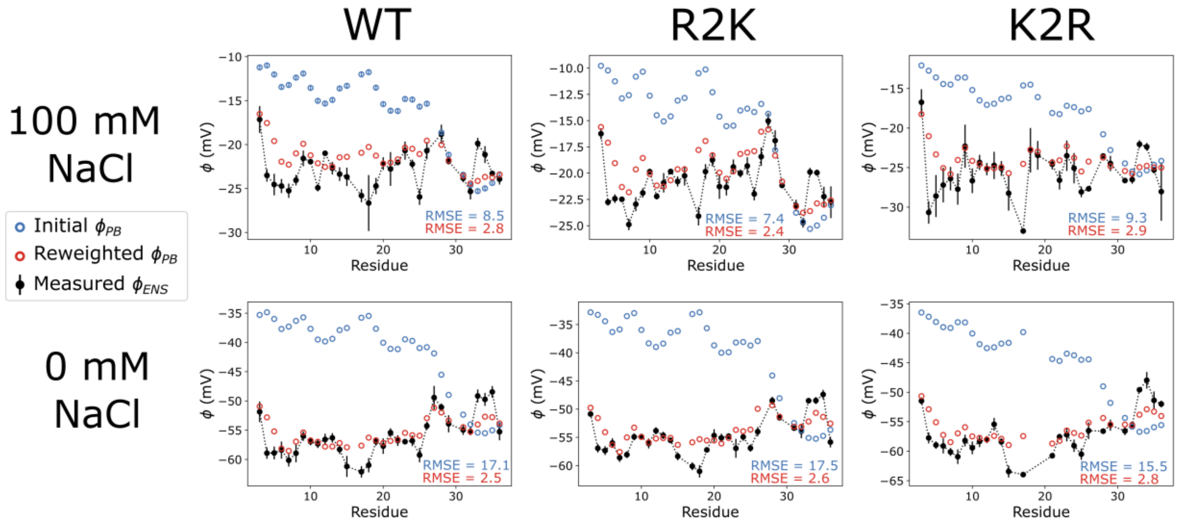


**Figure S4: Reweighting of WT (left, same as Figure 3b,c), R2K (middle), and K2R (right) ensembles at 100 mM (top) and 0 mM (bottom) NaCl**. Unweighted ensemble $\phi_{PB}$ values are shown in blue, while reweighted values are in red. Experimental $\phi_{ENS}$ measurements are in black, along with experimental errors (vertical lines). RMSE to $\phi_{ENS}$ (mV) is shown for both unweighted and reweighted $\phi_{PB}$ values.

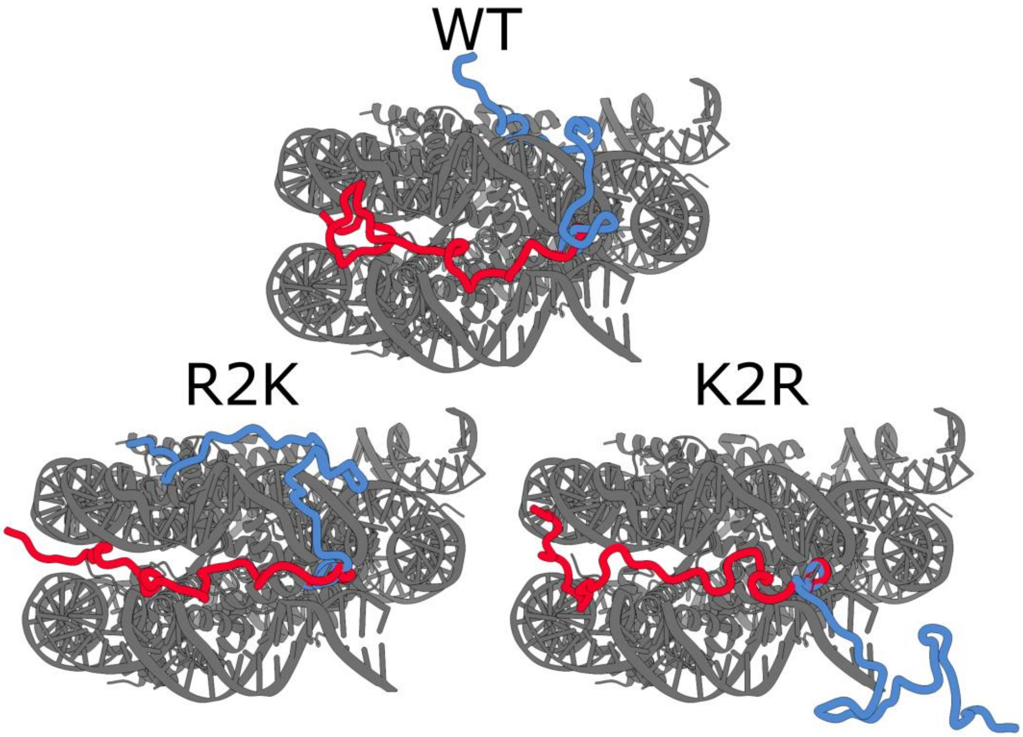


**Figure S5: NCP structures showing tails at $\phi_{PB,tail}$ extremes for each ensemble.** NCP structure with H3 tails with most negative (red) and least negative (blue) $\phi_{PB,tail}$ values for the WT (top, same as Figure 4a), R2K (bottom, left) and K2R (bottom, right) ensembles (100 mM NaCl).

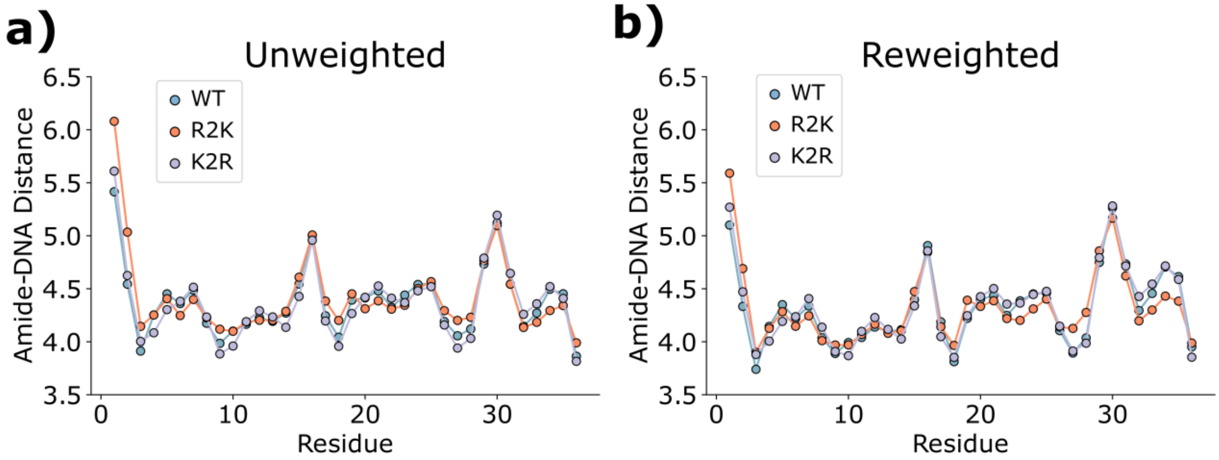


**Figure S6: Ensemble average tail-DNA distances for unweighted and reweighted ensembles.** Ensemble average distance from a given amide nitrogen to the nearest heavy atom within the nucleosomal DNA for the WT (blue), R2K (orange), and K2R (lavender) H3 tails from unweighted (a) and weighted (b) ensembles (100 mM NaCl).

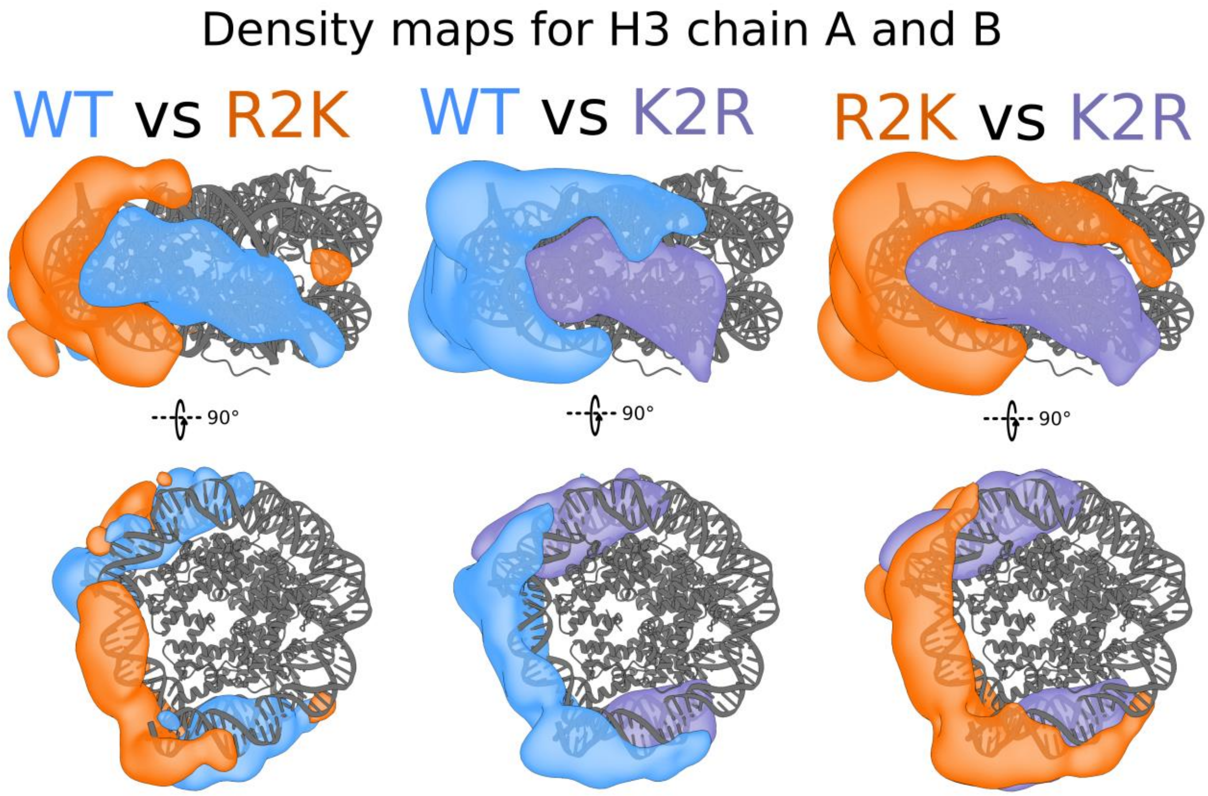


**Figure S7: Density maps including both H3 'A' and 'B'.** Difference density maps for high salt reweighted ensembles, as Figure 4d-f, but where densities are shown for all backbone heavy atoms in *both* tails 'A' and 'B'. Contour levels are set to the same value as in Figure 4. To maintain identical contouring between figures, grids for these maps are assembled by first generating a grid for 'A' and 'B' separately, then combining them, so the sum of all weights is equal to 2 rather than 1 as in Figure 4.

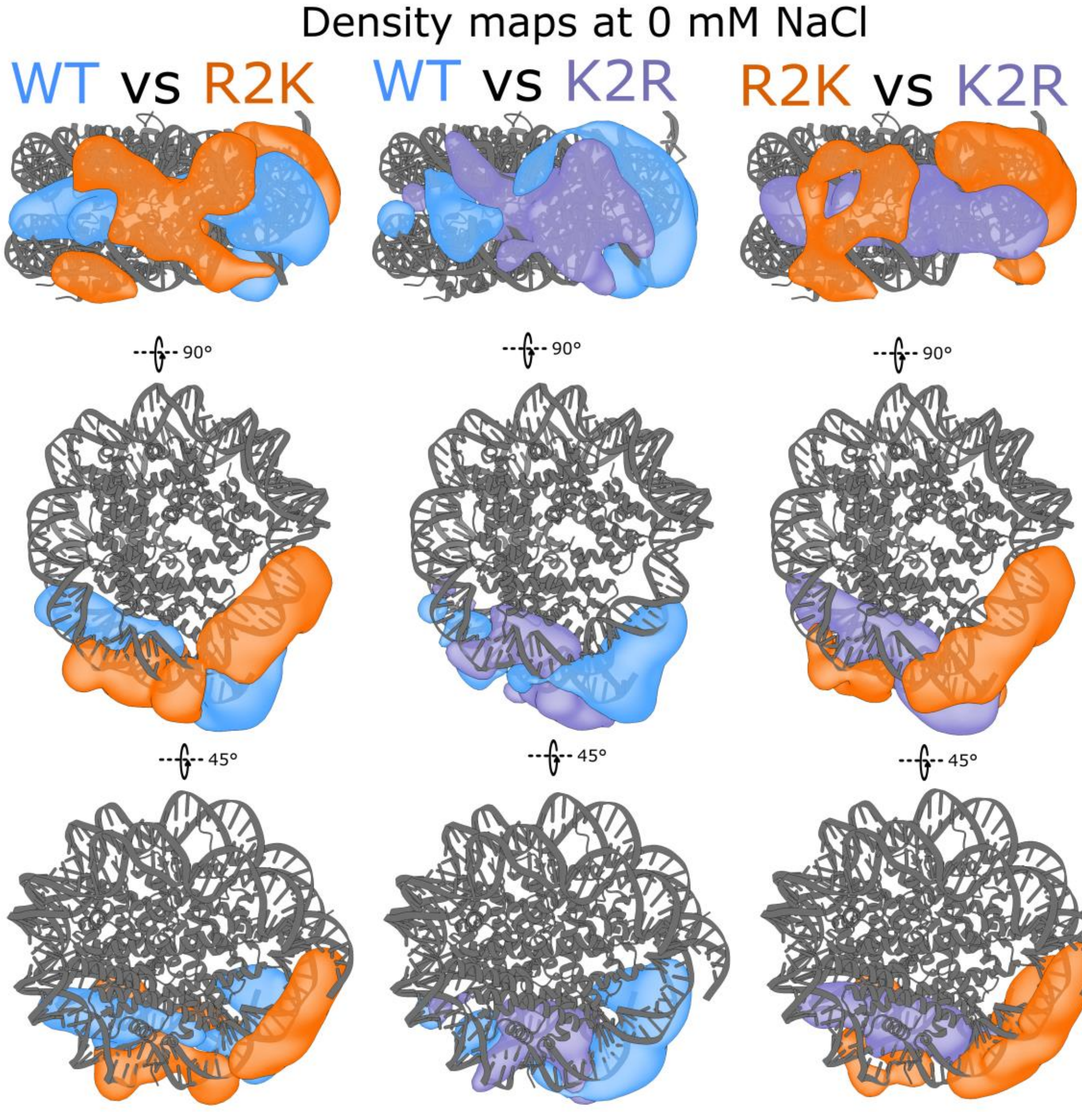


**Figure S8: Difference density maps of ensembles reweighted to data at 0 mM NaCl.** Difference maps comparing WT *vs* R2K (left), WT *vs* K2R (right), and R2K *vs* K2R (right) ensembles reweighted to $\phi_{ENS}$ measurements at 0 mM NaCl. Density for WT is shown in blue, R2K in orange, and K2R in lavender. Contour levels are set to the same value as in Figure 4.

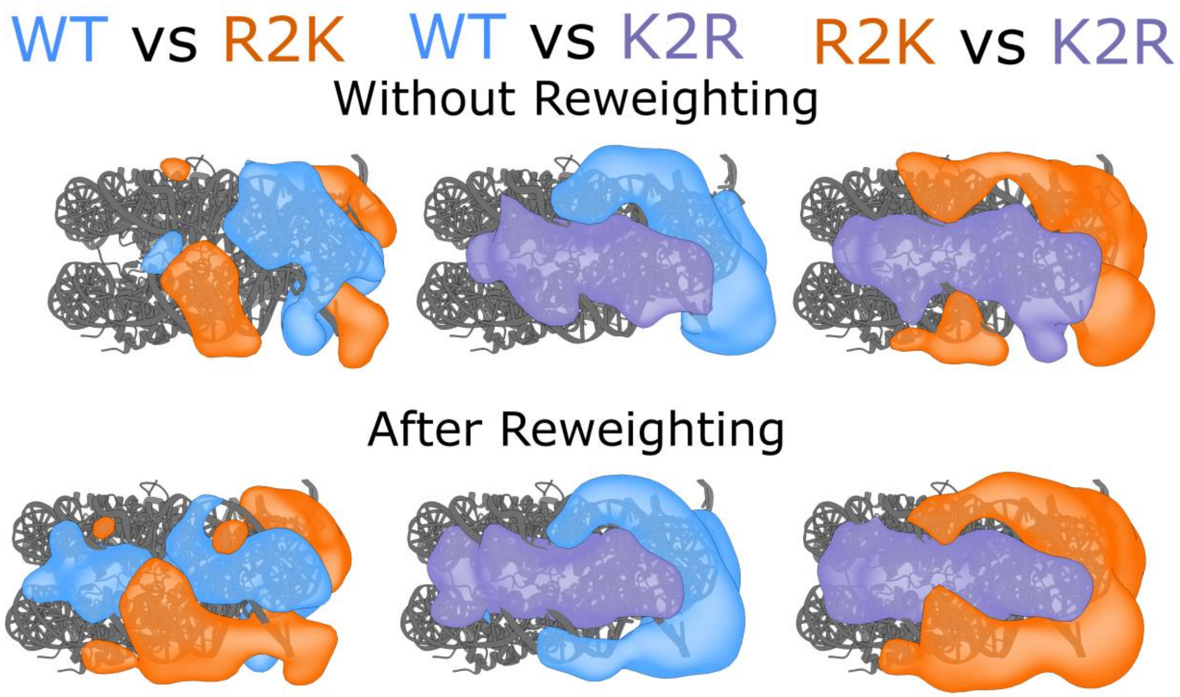


**Figure S9: Difference density maps before and after reweighting to $\phi_{ENS}$ measurements.** Difference maps comparing WT *vs* R2K (left), WT *vs* K2R (right), and R2K *vs* K2R (right). Density for WT is shown in blue, R2K in orange, and K2R in lavender. Contour levels are set to the same value as in Figure 4. Top row shows maps generated from unweighted ensembles, while bottom maps derive from ensembles reweighted to $\phi_{ENS}$ measurements at 100 mM NaCl (Figure 4d-f). Density differences are apparent in some cases prior to reweighting, but are accentuated by the reweighting process, especially in the case of the WT and R2K ensembles.

**Captions for supporting movies (movies are attached as separate files):**

**Supporting Movie S1: Movie of contours for R2K, WT, and K2R reweighted ensembles.** Videos move through density contour levels from high to low, starting at 0.05% and ending at 0.001%. The area of highest probability density (and therefore highest contouring) is along the side of the nucleosome, between DNA gyres, and this density is highest in the K2R variant, followed by WT, then R2K. Only at lower contour levels does probability density begin to appear around the edges of the NCP.

**Supporting Movies S2 and S3: Movie of contours from side (S2) and top (S3) view of density grids for R2K – K2R difference map.** Videos move through density contour levels from high to low, starting at 0.02% and ending at 0.0003%. It is difficult to visualize the probability density in 3D space, and scrolling through contours allows visualization of how the densities for R2K and K2R variants differ. Specifically, the K2R variant has high probability density in a relatively narrow space along the DNA gyres, which therefore appears at high contouring. In comparison, the R2K variant density is diffuse, and only begins to emerge at lower contour levels, but covers a larger area once it appears.
